# An integrated single-cell atlas of peripheral glia reveals progenitor-like states and enables cross-study mapping of dorsal root ganglia

**DOI:** 10.64898/2026.01.29.702614

**Authors:** Pauline Meriau, Michael B. Thomsen, Oshri Avraham, Brent Trauterman, Valeria Cavalli

**Author notes:** Correspondence to, Valeria Cavalli, Department of Neuroscience, Washington University School of Medicine, MSC 8108-96-07, 660 South Euclid Avenue, St. Louis, MO 63110-1093. co-first authors (Pauline Meriau, Michael B. Thomsen). Department of Cellular Biology, University of Georgia, Athens, GA 30602.

## Abstract

Sensory ganglia in the peripheral nervous system contain diverse glial populations that modulate sensory signaling, respond to injury and contribute to tissue homeostasis. Satellite glial cells (SGCs) surrounding neuronal soma in dorsal root ganglia (DRG) were suggested to retain developmental potential, but the identity of progenitor cells remains undefined. To capture glial diversity, we assembled a comprehensive single-cell transcriptomic atlas by integrating over 200,000 DRG and sciatic nerve transcriptomes across multiple studies and injury paradigms. High-resolution clustering resolved 28 cell types and demonstrated significant transcriptional heterogeneity within SGCs and Schwann cells, including repair and reactive sub-states. We identified two distinct populations of progenitor cells that reflect different states in the progenitor trajectory. Functionally, progenitor cell numbers increase after injury, and endothelin signaling regulates glial cell proliferation early in development. This integrated DRG and peripheral nerve atlas represents an essential resource for exploring new features of the peripheral nervous system.

## Introduction

The peripheral nervous system (PNS) relies on specialized glial populations to maintain neuronal function, modulate sensory signaling, and coordinate responses to injury. Within dorsal root ganglia (DRG), satellite glial cells (SGCs) and Schwann cells (SCs) support neuronal homeostasis while adapting to physiological and pathological stimuli. Emerging evidence suggests that these glial cells exhibit substantial functional diversity ^1, 2, 3^. However, a unified and high-resolution molecular framework that captures the full spectrum of peripheral glial states across conditions has remained lacking. Resolving this complexity is essential for understanding how glial cells contribute to sensory function, nerve regeneration, pain and disease.

The soma of each sensory neuron is enveloped by multiple satellite glial cells (SGCs), forming an intimate anatomical and functional unit ^4, 5, 6^. While once viewed as passive support cells, SGCs are now recognized as active regulators of neuronal function by modulating the perisomatic microenvironment ^7, 8, 9^. SGCs thus regulate the transmission of sensory inputs such as pain, itch, touch, pressure, and bodily position to the central nervous system, informing the brain on the internal and external state of the body. SGCs respond to various types of nerve injuries, positioning them as key contributors to pain in multiple conditions, sensory dysfunctions and nerve repair ^7, 8, 9^.

Another way SGCs can assist with nerve repair is via their stemness potential. Adult neurons lost to injury were suggested to be replaced by newly born ones deriving from a pool of DRG stem cells ^10, 11, 12, 13, 14^. Recent evidence suggests that these stem cells are SGCs. We now know that adult SGCs retain expression of genes related to glial progenitors and pluripotency, such as *Nestin*, *Sox2* and *Sox10* ^1, 15, 16, 17^. Additionally, lineage tracing experiments showed that Nestin-positive SGCs can generate new SGCs under homeostatic conditions, or sensory neurons after nerve injury ^17^. Adult SGCs/DRG stem cells can give rise to neurons, other glial cells or pericytes, suggesting that their multipotency and cell fate is modulated by environmental cues ^16, 17, 18, 19^. However, the niche in which these SGCs/DRG stem cells reside has not been clearly established. A recent study indicates that promyelinating SCs are nested in the convoluted initial axon segment near the cell soma, known as the Cajal’s initial glomerulus (IG) ^20^. Whether Cajal’s IG also contains SGCs/DRG stem cells and thus represents a unique glial developmental microdomain in the DRG remains unclear. Furthermore, the detailed molecular profile of these SGCs/DRG stem cells and the signaling mechanisms that regulate their proliferation or differentiation remain unknown.

Single-cell transcriptomics has transformed our ability to resolve cellular diversity in complex tissues, including the peripheral nervous system. Recent studies have begun to characterize glial populations in dorsal root ganglia and peripheral nerves, identifying major cell types and injury-associated states. However, these analyses have largely been performed in isolation, often using distinct experimental platforms, sequencing modalities, and annotation strategies, resulting in inconsistencies in cell-type definitions and limited resolution of rare or transitional states. Most studies have described SGCs and SCs but did not differentiate between myelinating SCs (mSCs) and non-myelinating (nmSCs) ^1, 2, 21, 22^. Other studies in sympathetic ganglia classified all glial cells as SCs ^23^. Only few studies have described all three types of glial cells in the DRG ^3, 24^. This apparent discrepancy may be due to the lack of precise molecular markers that differentiate nmSCs from SGC subtypes, sequencing depth, or differences in tissue dissection techniques. As a result, determining whether distinct SGC subtypes exist, including a potential progenitor population, has remained a challenge.

Here, we generated a comprehensive and unified single-cell and single-nucleus atlas of mouse DRG and sciatic nerve by integrating over 200,000 across multiple studies, developmental stages, and injury paradigms that resolves the transcriptional heterogeneity of peripheral glia. We reveal that SGCs present as a relatively uniform cell type at the transcriptional level, with only immune-responsive and immediate early gene-expressing states distinguishable from the major SGC population. Despite the lack of distinguishing features at the transcriptional level, SGCs exhibit functional specialization at the protein level that correlates with the neuronal subtype they envelop. We identify rare progenitor-like progenitor populations with proliferative and immature transcriptional signatures, localized within the Cajal’s IG. Using complementary computational approaches, including trajectory inference and RNA velocity, we reveal that the progenitor-like cells have the potential to generate glial cells in both myelinating and non-myelinating lineage. Both SGCs and progenitor cells express the Endothelin B receptor (ETBR), and we demonstrate that endothelin signaling via ETBR drives SGC proliferation in embryonic and early post-natal developmental stages. Together, these findings uncover the cellular and molecular complexity of peripheral glia and establish a unified reference framework for peripheral glia, setting the stage for new discoveries into their roles in peripheral nervous system function.

## Results

### An integrated single cell atlas of the mouse DRG and sciatic nerve

To construct a comprehensive single cell reference atlas of the DRG microenvironment, we integrated publicly available ^1, 21, 22, 25, 26, 27, 28, 29, 30, 31, 32, 33^ and newly generated single-cell and single-nucleus transcriptome datasets. We chose to integrate datasets from mouse (DRG) and sciatic nerves (SN) to better unify glial cell annotation, since DRG contains neuronal soma, SGCs, axon bundles, and associated Schwann cells (SCs) that extend into the nerve, whereas SN lacks SGCs but contains an abundance of SCs (Figure 1A). Our integrated data set comprises a total of 136,549 DRG cells and 90,972 SN cells (Figures 1B; S1). These datasets originated from diverse technologies (cells vs. nuclei; Drop-seq vs. 10X Genomics), multiple biological conditions (four different injury paradigms plus uninjured controls), different developmental time points (p1 and adult) and varied experimental designs (Figures 1C, Table S1). To account for technical differences across datasets, we used a robust integration pipeline to mitigate batch effects and highlight genuine biological differences among cell types and tissues (see Methods). Integration markedly improved mixing of cells across studies, sequencing preparations, tissues, and technologies while preserving biological cluster structure (Figure S2). Following integration, we first classified cells based on their principal cell type (Class), including neurons *(Snap25, Syt1),* glial cells (*Sox10*, *Plp1*), fibroblasts (*Col6a1, Col1a1*), endothelial cells (*Pecam1, Flt1*), immune cells (*Aif1, Ptprc*), mural cells (*Rgs5, Acta2*), and erythrocytes (*Hba-a1*) (Figure S3). We then iteratively subset, clustered, and annotated each cell type to identify a total of 7 classes, 28 major cell types, and 84 minor cell types (Figures 1B-D; S4, Movie S1). This hierarchical clustering approach allowed us to characterize the full scope of cell diversity harmonized between both DRG and SN compartments. Importantly, clusters were well represented across all studies and conditions, indicating successful integration without artificial segregation by batch or metadata category (Figure 1C, E; Table S1). In line with their known anatomical organization (Figure 1A), SN datasets exhibited abundant SCs (both mSCs and nmSCs) and, as expected, lacked sensory neurons (Neuron) and SGCs (Figure 1E). DRG datasets, by contrast, contained robust populations of Neurons and SGCs (Figure 1E), as well as all other cells present in the nerve, although in overall lower proportion. Established cell type-specific marker genes were consistently enriched in both DRG and SN, and our analysis defined distinctive gene expression signatures for each of the 28 major and 84 minor cell types identified in our dataset (Figure 1D, Table S2).

**Figure 1.**
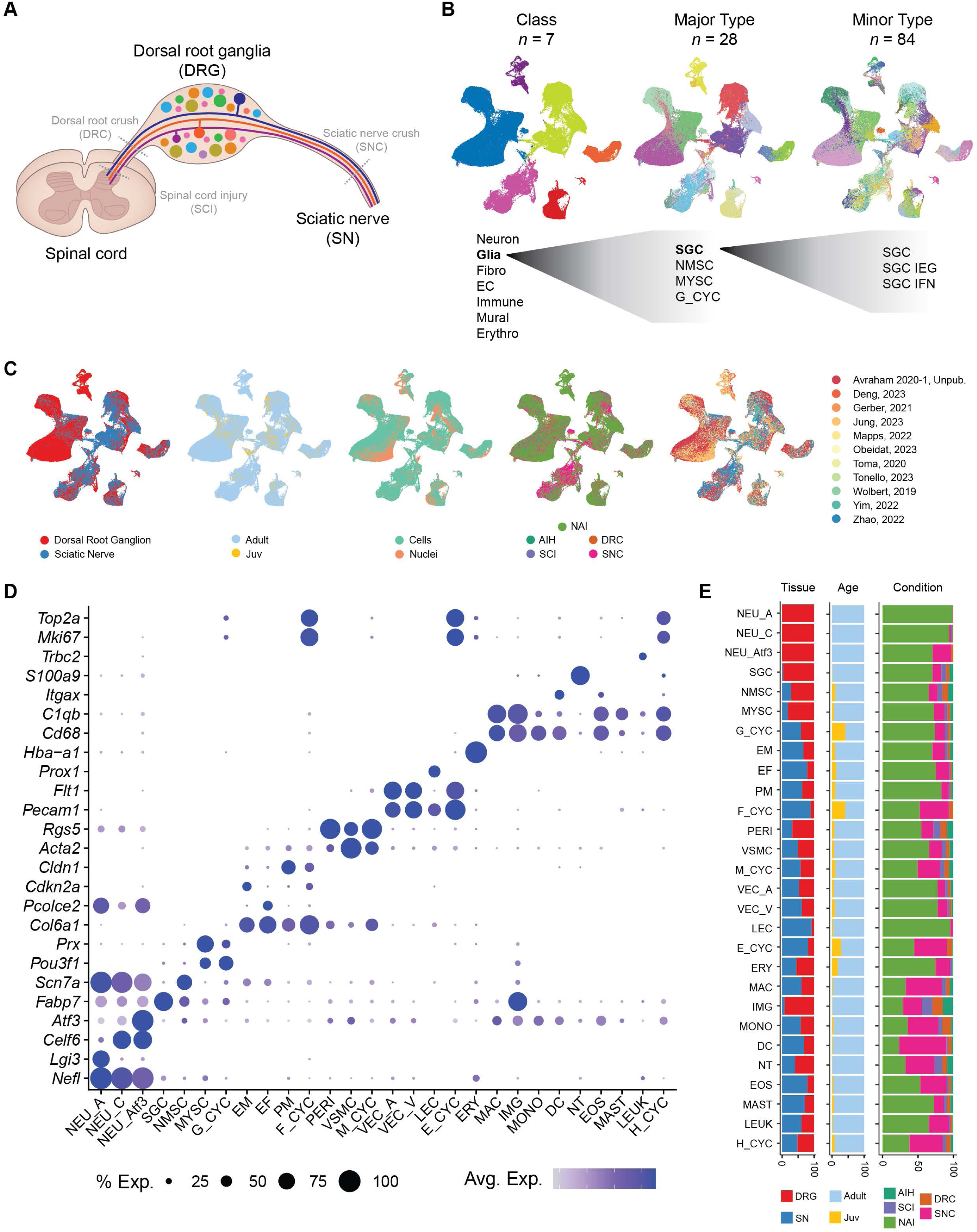
Integrated single-cell atlas of dorsal root ganglion and sciatic nerve. (**A**) Schematic of experimental design. (**B**) UMAP visualizations of the integrated dataset (136,549 DRG cells; 90,972 SN cells) colored by hierarchical clustering level: class (left), major type (middle), and minor type (right). Example cluster annotations are shown below each UMAP to illustrate the hierarchical relationship between clustering levels. (**C**) UMAP visualizations colored by metadata category: tissue of origin, animal age, sequencing preparation (single-cell or single-nucleus), treatment condition, and source study. (**D**) Dotplot of marker gene expression across major cell types. Dot size indicates percentage of cells expressing each gene; color intensity indicates average expression level. (NEU_A, A fiber neurons; NEU_C, C fiber neurons; NEU_Atf3, Atf3-positive neurons; SGC, satellite glial cells; NMSC, non-myelinating Schwann cells; MYSC, myelinating Schwann cells; G_CYC, mitotic glial cells; EM, endoneurial mesenchymal cells; EF, epineurial fibroblasts; PM, perineurial mesenchymal cells; F_CYC, mitotic fibroblasts; PERI, pericytes; VSMC, vascular smooth muscle cells; M_CYC, mitotic mural cells; VEC_A, arterial vascular endothelial cells; VEC_V, veinous vascular endothelial cells; LEC, lymphatic endothelial cells; E_CYC, mitotic endothelial cells; ERY, erythrocytes; MAC, macrophages; IMG, imoonglia; MONO, monocytes; DC, dendritic cells; NT, neutrophils; EOS, eosinophils; MAST, mast cells; LEUK, leukocytes; H_CYC, mitotic hematopoietic cells). (**E**) Stacked bar graphs depicting the relative proportion of cells within each major cell type by tissue, age, and treatment condition.

This resource provides unprecedented resolution of PNS cell diversity and establishes a validated cross-study reference for the field. Notably, this integrated atlas provided an ideal launching point for in-depth annotation and analysis of glial cells, whose transcriptional diversity and context-specific roles are increasingly recognized as pivotal in PNS function, maintenance and repair ^7, 8, 9, 34, 35^. We therefore next leveraged the atlas to dissect glial cell heterogeneity in naïve and injured contexts.

#### Diversity of peripheral glial cells

Annotation of major glial cell types in single cell datasets from peripheral ganglia has been challenging. Studies in the DRG in multiple species have identified SGCs, mSCs, and nmSCs ^3, 24^, while other studies in mice describe SGCs and SCs, but did not identify nmSCs ^1, 21, 22, 25^. Other studies in sympathetic ganglia classified all glial cells as SCs ^23^. Because our integrated data set includes both nerve and DRG, glial annotation can be precisely harmonized. We first used marker gene enrichment to define each major glia cluster, labeled by expression of *Plp1*, which identified well-characterized marker genes such as *Fabp7* in SGCs ^21, 25^, *Scn7a* in nmSCs ^3, 36^ and *Mbp* in mSCs ^37^ (Figure 2A-D). Immunolabeling in DRG tissue highlighted the distinct spatial distributions of each major glial cell type, with SGCs forming the typical circular pattern in neuron soma rich areas and nmSCs and mSCs primarily occupying the axon rich region within the DRG (Figure 2E). On rare occasion Scn7a-positive cells extend thin processes into the SGCs rich area (Figure 2E), as recently described ^2^ .

**Figure 2.**
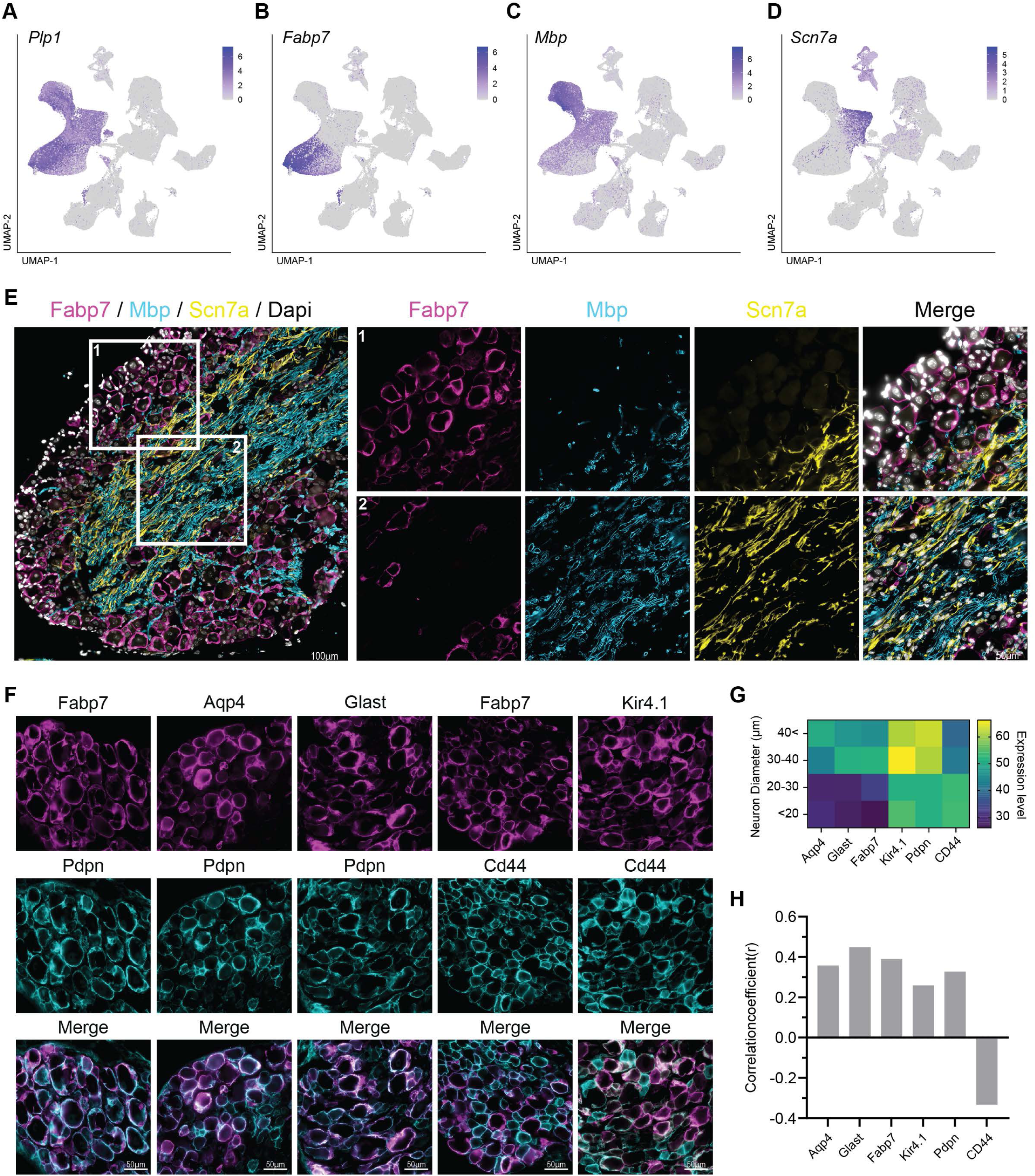
Molecular characterization of peripheral glial populations. (**A-D**) UMAP feature plots showing expression of glial marker genes: *Plp1* (**A**), *Fabp7* (**B**), *Mbp* (**C**), and *Scn7a* (**D**). (**E**) Representative immunofluorescence images of neuron soma dense area (1) and axon bundle dense area (2) of DRG cross-section labeled for Fabp7 (magenta) enriched in SGCs, Mbp (cyan), enriched in mSC, Scn7a (yellow) enriched in nmSC and Dapi (grey). Scale bars: 100 µm (overview), 50 µm (high magnification). (**F**) Representative immunofluorescence images of SGC markers; Fabp7, Aqp4, Glast, Pdpn, Cd44 and Kir4.1 expression and colocalization. Scale bars: 50 µm. (**G**) Heatmap of SGCs markers expression level (mean gray value); Fabp7, Aqp4, Glast, Pdpn, Cd44 and Kir4.1 binned according to the size of the associated neuron. (**H**) Pearson correlation coefficients of Fabp7, Aqp4, Glast, Kir4.1, Pdpn expression showing positive correlations with neuron size and a negative correlation with Cd44.

We next validated the expression of novel SGC markers using a combination of transcript-level visualization in UMAP space and immunofluorescence (Figure 2F, S5). Podoplanin (Pdpn) and Cd44 emerged as two newly identified SGC protein markers (Figure 2F, S5). While transcripts for Pdpn and Cd44 were detected in less than 50% of SGCs, their robust protein expression in SGCs was confirmed by co-localization with Fabp7 and Sox10 (Figures 2F, S5A-C, G). Although at the transcriptomic level, *Pdpn* is also expressed in fibroblasts (Figure S5A), immunostaining showed very low co-localization of Pdpn with the fibroblast marker Decorin (Dcn) compared to established SGC markers, confirming that Pdpn specifically labels SGCs (Figure S5D-F). Cd44 transcripts were detected in both Sox10 SGCs and nmSCs (Figure S5A), and the Cd44 protein was observed in both SGCs as well as nmSCs, which are enriched in the axon rich area of the DRG (Figure S5G-I).

We then examined if these newly identified SGC makers underlie SGC specialization around the neuron they envelop. Although SGCs represent a relatively unified population at the transcriptional level (Figure 2B), we observed that the protein expression levels of Fabp7, Pdpn, Glast, Aqp4, and Kir4.1 were higher in SGCs enveloping larger neuronal soma (Figures 2F-H, S5J). In contrast, Cd44 displayed an inverse relationship, showing higher expression level in SGCs around smaller neurons (Figures 2F-H, S5H-J). These size-dependent expression patterns suggest that SGC specialization is achieved through protein-level adjustments to meet the specific needs of their target neuron.

#### Transcriptional diversity of minor glial cell types

To resolve the full transcriptional heterogeneity of peripheral glia, we then focused on the minor cell type clusters of glia within our hierarchical clustering framework (Figure 1B). Our analysis identified 11 distinct glial subpopulations, each defined by a characteristic gene expression profile (Figures 3A, S6B, Movie S2, Table S3). When we compared DRG-versus SN-derived glia, the same 11 clusters were identifiable in both tissues, albeit at different proportions (Figure 3A-C, Figure S6A, C). As expected, SGCs predominated in the DRG, whereas nmSCs and mSCs were more prevalent in the SN (Figure 3B). Integrating data from >35,000 DRG-derived SGCs identified three subpopulations: a homeostatic population (SGC), immune-responsive SGCs (SGC IFN), and immediate early gene-expressing SGCs (SGC IEG). SGC IFN cells are enriched for interferon-induced transcripts including *Ifit1*, *Ifit3a*, *Gbp2*, and *Stat1* ^2, 21^. SGC IFN likely represent a population that respond to viral infection, as recently demonstrated for Herpes Simplex Virus ^2^, supporting potential involvement in antiviral protection ^8, 38^. SGC IEG cells express classical IEGs such as *Fos* and *Jun* and may represent an active or poised state to respond dynamically to normal neuronal signaling or pathological conditions. Alternatively, this population may represent a transient state induced by tissue dissociation.

**Figure 3.**
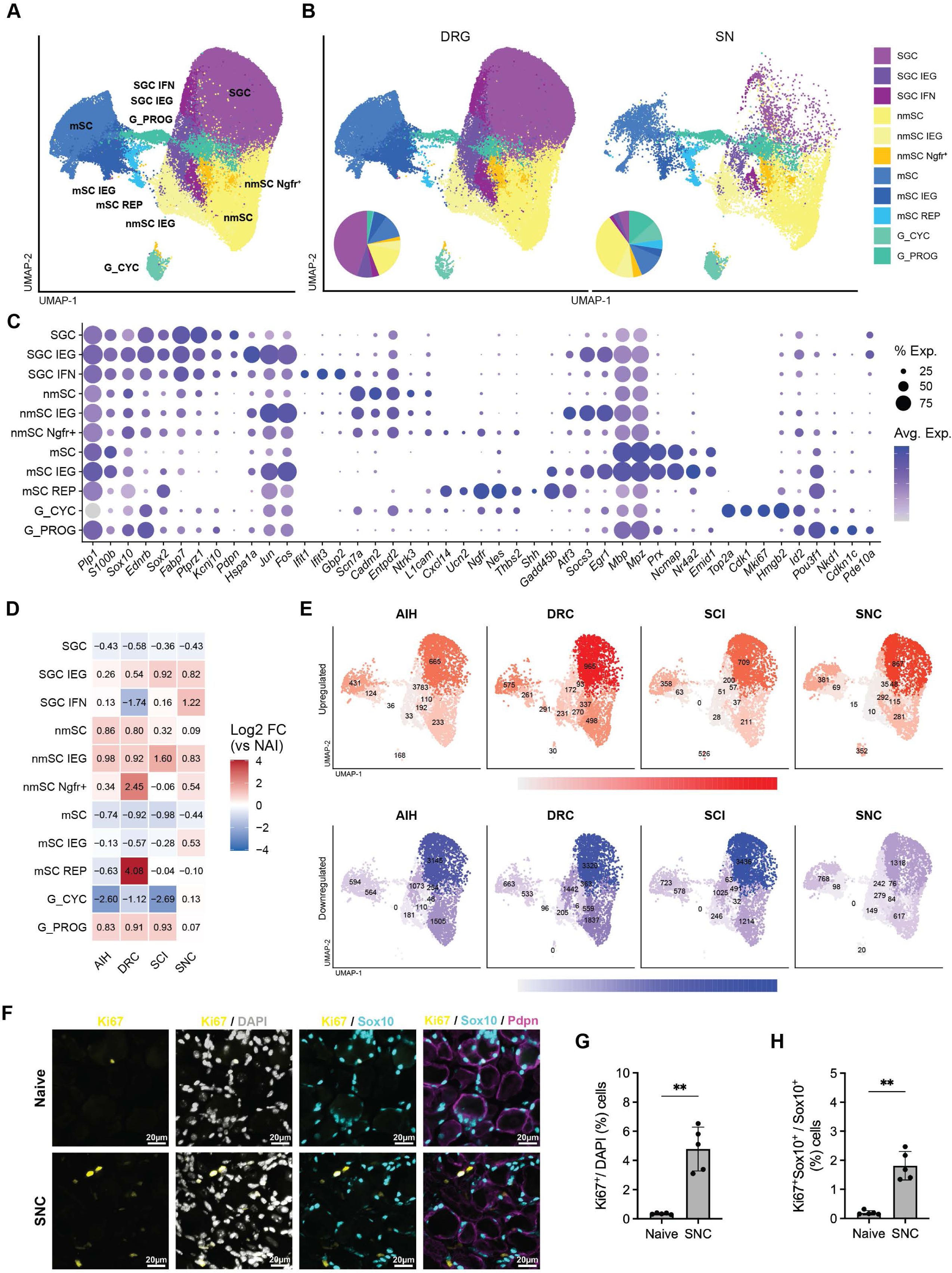
Peripheral glial subpopulations and transcriptional responses to injury. (**A**) UMAP visualization of 11 glial subpopulations identified by minor cell type clustering (SGC, satellite glial cell; SGC IFN, interferon-responsive SGC; SGC IEG, immediate early gene-expressing SGC; nmSC, non-myelinating Schwann cell; nmSC IEG, immediate early gene-expressing nmSC; nmSC Ngfr+, Ngfr-positive nmSC; mSC, myelinating Schwann cell; mSC IEG, immediate early gene-expressing mSC; mSC REP, repair mSC; G_CYC, mitotic glial cells; G_PROG, post-mitotic progenitors). (**B**) UMAP visualization split by tissue (DRG, left; SN, right). Inset pie charts show relative abundance of each glial subpopulation per tissue. (**C**) Dotplot of marker gene expression across glial subpopulations. Dot size indicates percentage of cells expressing each gene, color intensity indicates average expression level. (**D**) Heatmap of log2 fold change in glial subpopulation abundance relative to naïve (NAI) across four injury conditions: acute intermittent hypoxia (AIH), dorsal root crush (DRC), spinal cord injury (SCI), and sciatic nerve crush (SNC). (**E**) UMAP visualizations of DRG cells showing the number of significantly differentially expressed genes (adjusted p < 0.05) per subpopulation following each injury condition. Top row, upregulated genes; bottom row, downregulated genes. Each panel displays only cells from the indicated injury condition. (**F**) Representative immunofluorescence images of DRG sections from naïve and SNC mice stained for Ki67 (yellow), SOX10 (cyan), Pdpn (magenta), and DAPI (white). Scale bars, 20 μm. (**G**) Quantification of Ki67⁺ cells expressed as a percentage of total cells in DRG sections from naïve and SNC mice. (**H**) Quantification of Ki67⁺SOX10⁺ cells expressed as a percentage of total SOX10⁺ cells. SNC injury significantly increased both the overall number of proliferating cells and the proportion of proliferating SOX10⁺ glial cells in the DRG. Data are presented as mean ± s.e.m.; each dot represents one biological replicate. Statistical significance was determined using an unpaired two-tailed Student’s t-test. P < 0.01 (**).

SCs similarly resolved into distinct subpopulations. Both mSCs and nmSCs included homeostatic and IEG-expressing clusters, paralleling the pattern observed in SGCs (Figure 3A-C). Additionally, we identified injury-associated populations in both SCs types. mSC REP cells, marked by expression of the repair SCs marker gene *Shh* ^39^, were largely derived from nerve datasets and were most abundant following dorsal root crush (DRC) in DRG samples (Figure 3D-E). nmSC Ngfr+ cells exhibited markers of repair SCs, including *Ngfr* and *Nes*, but lacked *Shh* expression (Figure 3C). These cells were enriched following DRC and sciatic nerve crush (SNC) in the DRG (Figure 3E), and their transcriptional profile resembles the highly ramified glia described at the morphological level surrounding proximal processes of large-diameter DRG neurons ^40^.

Our analysis also identified two progenitor-like populations, G_CYC and G_PROG (Figure 3A-C). G_CYC cells were defined by robust expression of proliferation markers, including *Mki67*, *Top2a*, *Plk1*, *Cdk1*, and GO enrichment confirmed strong representation of mitotic processes such as DNA replication, chromosome segregation, and nuclear division (Figure 3C, S6D). In contrast, G_PROG cells expressed markers associated with immature glia, including *Pou3f1*, *Id2,* and *Cdkn1c*, but lacked signatures of active cell division (Figure 3C).

To complement cluster-based analyses, we applied weighted gene co-expression network analysis (WGCNA), identifying 14 co-expression modules with distinct cell type enrichment patterns (Figure S7A, S7E-F). Modules associated with mSCs reflected canonical myelination and axon ensheathment programs (Figure S7B), whereas nmSCs-enriched modules featured neuron projection guidance and cellular response pathways (Figure S7C). Progenitor-associated modules highlighted ribosome biogenesis and mRNA processing pathways. Notably, four modules showed selective enrichment in SGCs, encompassing developmental signaling pathways, lipid and cholesterol metabolism, and proteostatic maintenance (Figure S7D), underscoring functional specialization despite their relatively uniform transcriptional profile.

Together, these analyses provide a consolidated reference for peripheral glial populations, serving as a resource to address cross-study inconsistencies in annotation. To facilitate comparison with prior work, we mapped each cluster to previously reported glial cell annotations, highlighting correspondences and clarifying discrepancies (Table 1). To demonstrate the practical utility of this atlas as a community reference, we performed reference mapping and label transfer onto several independent datasets that were not part of its construction, including a Fragile X model (Fmr1-KO; Figure S8) ^41^, a sciatic nerve crush injury time-course (Figures S9, S10), and an aged DRG dataset (Figure S11) ^42^. In each case, label transfer assigned atlas-defined identities at high confidence and recovered the expected cell-type composition together with injury- or age-associated compositional shifts, supporting use of the atlas as a stable cross-study reference.

**Table 1.** Glia cluster nomenclature crosswalk. Mapping of the 11 glial cluster identities defined in this atlas (SGC, SGC IEG, SGC IFN, nmSC, nmSC IEG, nmSC Ngfr+, mSC, mSC IEG, mSC REP, G_CYC, G_PROG) against published annotations from prior single-cell studies of peripheral glia (Avraham 2020; Avraham 2021; Tonello 2023; Zhao 2022; Wolbert 2019; Jung 2023; Mapps 2022; Yim 2022; Toma 2020; Obeidat 2023; Gerber 2021). NP = cluster not present in the source dataset; NA = cluster not annotated by original authors. Asterisks (*) indicate that the original authors reported multiple distinct sub-clusters that were merged before annotation.

| Cell Type | Avraham, 2020 | Avraham, 2021 | Tonello, 2023 | Zhao, 2022 | Wolbert, 2019 | Jung, 2023 | Mapps, 2022 | Yim, 2022 | Toma, 2020 | Obeidat, 2023 | Gerber, 2021 |
| --- | --- | --- | --- | --- | --- | --- | --- | --- | --- | --- | --- |
| **SGC** | SGC | 'SGC 2',<br>'SGC 3' | 'SGCs I' | NP | NP | 'Satellite glia' | 'Sensory SGC' | NP | NP | 'SATG' | NP |
| **SGC IEG** | NA | NA | NA | NP | NP | NA | 'IEG SGC' | NP | NP | NA | NP |
| **SGC IFN** | NA | NA | NA | NP | NP | NA | 'IMM SGC' | NP | NP | NA | NP |
| **nmSC** | NA | 'SGC 4' | 'SGCs II' | 'nmSC' | 'mySC' | NA | 'Gen. Resident SGC' | 'Non-myelinating SCs' | Non-myel. SC, Uninjured | NA | 'nm(R)SC' |
| **nmSC IEG** | NA | NA | NA | NA | NA | NA | NA | NA | NA | NA | NA |
| **nmSC Ngfr+** | NA | NA | NA | NA | NA | NA | NA | NA | SC, Injured | NA | 'tSC' |
| **mSC** | SC | 'Schwann Cells' | 'Schwann Cells' | 'mSC' | 'mySC' | 'Myelinating Schwann' | 'Schwann cells' | 'Myelinating SCs' | Myel. SC, Uninjured | 'SCHW' | 'mSC'* |
| **mSC IEG** | NA | NA | NA | NA | NA | NA | NA | NA | NA | NA | 'mSC'* |
| **mSC REP** | NA | NA | NA | 'rSC' | NA | NA | NA | NA | SC, Injured | NA | NA |
| **G_CYC** | NA | NA | NA | 'prol.SC' | NA | NA | NA | NA | 'Prolif. SCs' | NA | 'prol. SC' |
| **G_PROG** | NA | 'SGC 1' | NA | 'prol.SC' | NA | NA | NA | NA | 'Prolif. SCs' | NA | 'pmSC' / 'iSC' |

NP - not present
NA - not annotated
\* Original authors noted multiple distinct clusters that were merged before annotation

#### Transcriptional responses of glial cells in multiple injury contexts

We next examined transcriptional changes in DRG glial populations across three paradigms of injury: SNC, DRC, spinal cord injury (SCI) and acute intermittent hypoxia (AIH). While AIH has been shown to elicit pro-regenerative responses in neurons ^43^, its effects on glial cells have not been characterized in detail. Across all conditions, SGCs exhibited the largest number of differentially expressed genes relative to other glial populations, consistent with their role as primary non-neuronal responders to perturbation in the DRG (Figures 3E, S12). Glial transcriptional responses broadly distinguished peripheral from central injury. SNC induced robust upregulation of injury-response genes in SGCs with comparatively modest downregulation, whereas central injuries (DRC and SCI) drove substantial transcript downregulation across multiple glial populations (Figure 3E). Jaccard similarity analysis confirmed this pattern: many glial subpopulations showed higher DEG concordance among central injuries and AIH than with SNC, suggesting a shared transcriptional program in response to central lesions and hypoxia that is distinct from peripheral nerve injury (Figure S12). Notably, AIH, despite lacking physical trauma, more closely resembled central injury profiles in both gene expression and cell abundance shifts. DRC elicited the most pronounced changes among the injury models, likely reflecting its anatomical proximity of the injury to the DRG. Both mSC REP and nmSC Ngfr+ populations were substantially expanded following DRC relative to other conditions, accompanied by heightened differential expression in these repair-associated populations (Figures 3D-E, S12). Progenitor populations showed a striking divergence in their injury responses: cycling progenitors (G_CYC) expanded selectively following SNC but showed minimal response to central injuries. G_CYC expansion was confirmed by immunofluorescence, which revealed a significant increase in Ki67⁺Sox10⁺ cells in the DRG following SNC injury (Figure 3 F-H). Post-mitotic progenitors (G_PROG), on the other hand, displayed the opposite pattern, responding more strongly to central injuries than to SNC (Figures 3D, E, S12). This differential mobilization suggests that distinct injury contexts may engage separate progenitor pools within the DRG.

#### A progenitor-like cell population nested in Cajal’s initial glomeruli in adult DRG

We and others previously identified a *Pou3f1*-positive SGC subtype ^1, 2, 3^, but low cell numbers prevented further detailed characterization. Because *Pou3f1* positive G_PROG cells appeared to encompass features of all three major glial types, we employed pseudotime trajectory analysis to delineate potential differentiation paths. This revealed a single trajectory axis running through the G_PROG cluster, with endpoints defined by gene signatures characteristic of either mSCs or nmSCs/SGCs (Figure 4A-D). This trajectory was robust to the inference method: Monocle3 pseudotime closely recapitulated the Slingshot ordering (gene-wise Spearman ρ = 0.94; Figure S13), and RNA velocity was consistent with transitions out of G_PROG toward both myelinating and non-myelinating fates. Velocity magnitude, a measure of transcriptional dynamism, increased across glial populations in all injury paradigms (SNC, DRC, SCI, AIH) relative to the naïve state, and the genes underlying this increase were resolved by a velocity-driver gene heatmap (Figure S14). Genes correlated with pseudotime showed clear polarity: canonical myelinating markers (*Mpz, Mbp, Pmp22, Emid1, Mag*) were associated with one end of the trajectory, whereas non-myelinating/SGC genes (*Dbi, Apoe, Ncam1, L1cam, Ednrb, Ptn*) defined the opposite end (Figure 4C-E). This analysis suggests that the G_PROG population exhibits a progenitor-like state, with transcriptional features compatible with generating either myelinating or non-myelinating glia.

**Figure 4.**
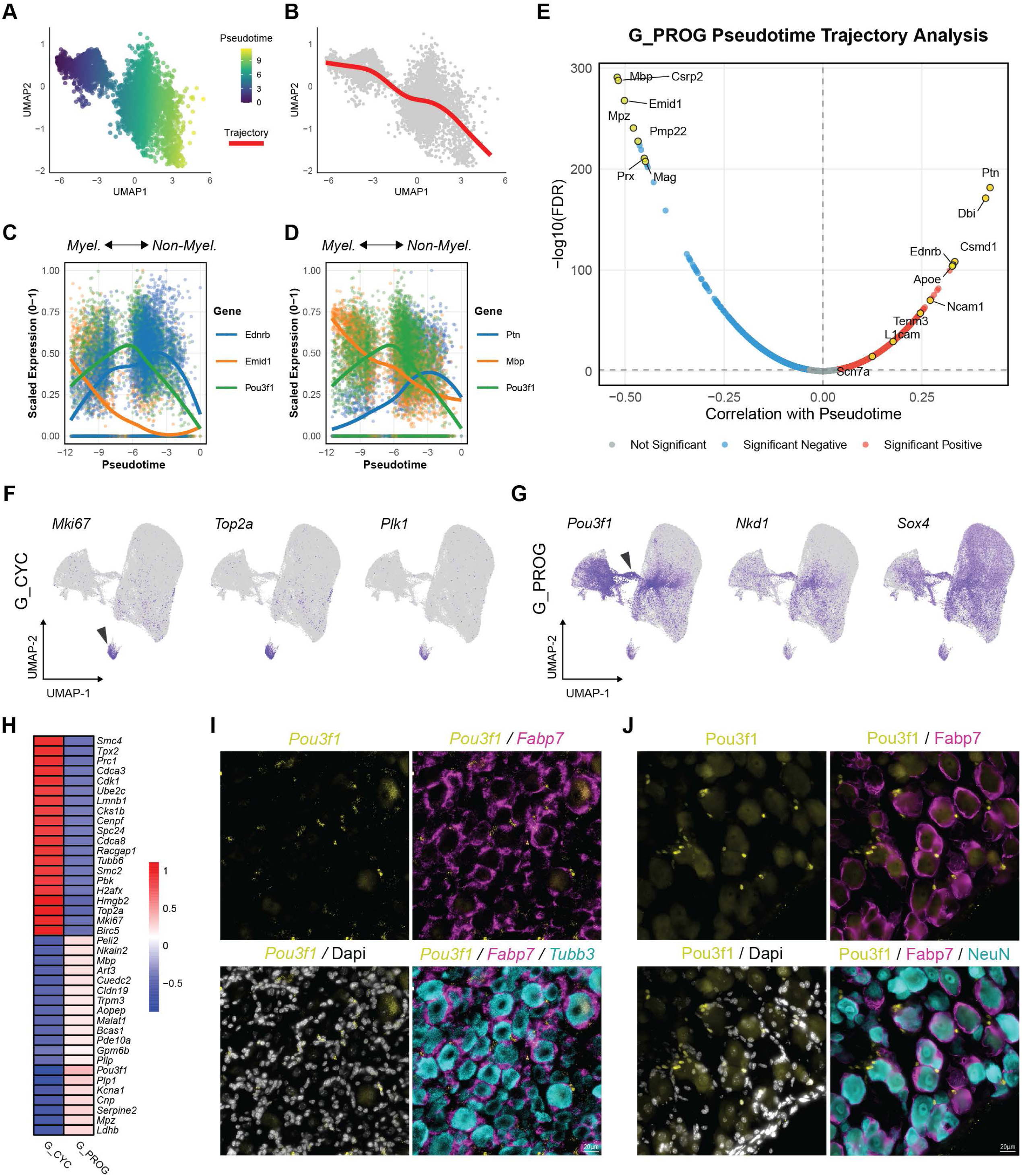
G_PROG cells define a progenitor-like glial state with myelinating and non-myelinating potential. (**A**) UMAP embedding of the G_PROG population colored by pseudotime progression, illustrating a continuous trajectory. (**B**) Inferred pseudotime trajectory overlaid on the UMAP embedding (red line), indicating the direction of lineage progression from progenitor to more differentiated states. (**C**) Gene expression dynamics along pseudotime for myelinating-associated (*Ermd1*), non-myelinating-associated (*Ednrb*) and progenitor-associated (*Pou3f1*) genes, illustrating transitions from progenitor to myelinating or non-myelinating states. (**D**) Gene expression dynamics along pseudotime for myelinating-associated (*Mbp*), non-myelinating-associated (*Ptn*) and progenitor-associated (*Pou3f1*) genes, illustrating transitions from progenitor to myelinating or non-myelinating states. (**E**) Volcano plot showing correlation between gene expression and pseudotime progression within the G_PROG II population. Genes with significant myelinating-associated fate (red) or non-myelinating-associated fate (blue) correlations (adjusted p < 0.05); representative markers of potential differentiation states are highlighted. (**F**) UMAP feature plots showing expression of cell cycle-associated genes *Mki67*, *Top2a*, and *Plk1*, identifying a small proliferative glial cell population (G_CYC; arrowhead). (**G**) UMAP feature plots showing expression of progenitor-associated genes *Pou3f1*, *Nkd1*, and *Sox4*, defining a progenitor-like glial population (G_PROG; arrowhead). (H) Heatmap of differentially expressed genes between G_PROG and G_CYC populations. (**I-J**) Representative in situ hybridization (**I**) and immunofluorescence (**J**) images showing Pou3f1 (yellow) and Fabp7 SGCs (magenta), with DAPI (white) and Tubb3 / NeuN neurons (cyan). Pou3f1 cells are very close to, or even part of, the ring of Fabp7 SGC surrounding neuron. Scale bars: 20 µm.

To provide an anatomical description of G_PROG and confirm the presence of these progenitors in intact adult DRGs, we examined their spatial relationship in DRG sections. *In situ* hybridization (ISH) and IF for the G_PROG marker *Pou3f1/*Oct6 and the SGC marker *Fabp7* (Figure 4I-J) revealed Pou3f1 positive cells in proximity of neuronal soma, with only partial overlap with Fabp7, confirming that a discrete population of immature glia exists in uninjured adult DRGs. Using high-resolution 3D clearing imaging to label neuron soma and their axons with peripherin and SGCs with Fabp7, we observed that SGCs form a continuous cellular sheath that encapsulates both the neuronal soma and the convoluted initial axon segment near the cell soma, known as the Cajal’s initial glomerulus (IG) (Figure S15E; Movie S3), similar to what has been observed in rats ^20^. This finding suggests that the IG represents a conserved anatomical niche for progenitor-like cells across species.

#### ETBR signaling promotes DRG glial cells proliferation early in development

To identify potential extrinsic regulators of fate decision and proliferation of G_PROG, we examined transcripts enriched at the nmSC/SGC pole of the trajectory. *Dbi*, which encodes diazepam binding inhibitor, was prominent among these and is notably enriched in mature SGCs (Figure 4E) ^44^. However, the strong and selective expression of *Ednrb* in these cells (Figure 3C, 4E) singled out endothelin signaling as a particularly compelling candidate pathway to investigate further. The endothelin B receptor (ETBR), encoded by *Ednrb*, can repress myelination ^45^, consistent with its expression in non-myelinating glial cells (Figures 3C, S15D) and enrichment in the non-myelinating/SGC axis of the pseudotime trajectory (Figure 4E). In addition, endothelin signaling via ETBR has been linked to glial cell proliferation, promoting Schwann cell proliferation in the PNS ^46^ and radial glial cell maintenance and proliferation in the postnatal subventricular zone ^47^. During DRG embryonic development, *Ednrb* is expressed in a small proportion of undifferentiated cells that also express *Sox2, Sox10, Nestin and Foxd3* at embryonic day 11.5 ^48^, consistent with the onset of gliogenesis in the DRG ^49^, suggesting that ETBR signaling mediates glial proliferation in the DRG early in development and possibly later in the adult stage. To investigate the role of the endothelin pathway in glial proliferation, we first characterized the cellular composition of 13.5 embryonic DRG (eDRG) primary cultures (Figure S16A). Gene expression analysis confirmed the expression of neural crest (*Foxd3, Rest, Sox2*), neuronal (*Tubb3*) and glial lineage markers (*Erbb3, Fabp7, Sox10*) in eDRG cultures (Figure S16B). Importantly, *Ednrb* transcripts were also detected, confirming the presence of the receptor at the transcriptional level in these cultures (Figure S16B). To further validate expression of a functional ETBR receptor, we performed calcium imaging using the Fluo-4-AM indicator. IRL1620, a specific ETBR agonist, triggered a robust increase in intracellular Ca2+ fluorescence in eDRG cells (Figure 5A). Following 5 days of treatment with endothelin-1 (ET-1) or IRL1620 a striking morphological reorganization occurred, with the emergence of “sphere-like structures,” defined as dense, three-dimensional cellular aggregates, with a minimum diameter of 100µm (Figures 5B-C S16C). These structures contained mixed cellular composition, comprising both NeuN positive neurons and Sox10 positive progenitor/glial cells (Figure 5B). To determine if these sphere-like structures resulted from active cell division, we performed EdU incorporation assays, which revealed a significant increase in the number of EdU-positive cells following IRL1620 treatment (Figure 5E). EdU-positive nuclei co-localized with the glial/progenitor marker Sox10, whereas no co-localization was observed with the neuronal marker NeuN (Figure 5D), supporting that ETBR-mediated proliferation is restricted to the glial lineage. Consistent with this mitogenic effect, mRNA expression analysis showed that ET-1 and IRL1620 treatment significantly upregulated the expression of the cell cycle regulator *Cdk1*, progenitor marker *Nestin*, and glial cell markers *Sox10*, and *Erbb3* (Figure 5F). Collectively, these data indicate that activation of the endothelin pathway drives the expansion of a glial progenitor-like population within eDRG cultures.

**Figure 5.**
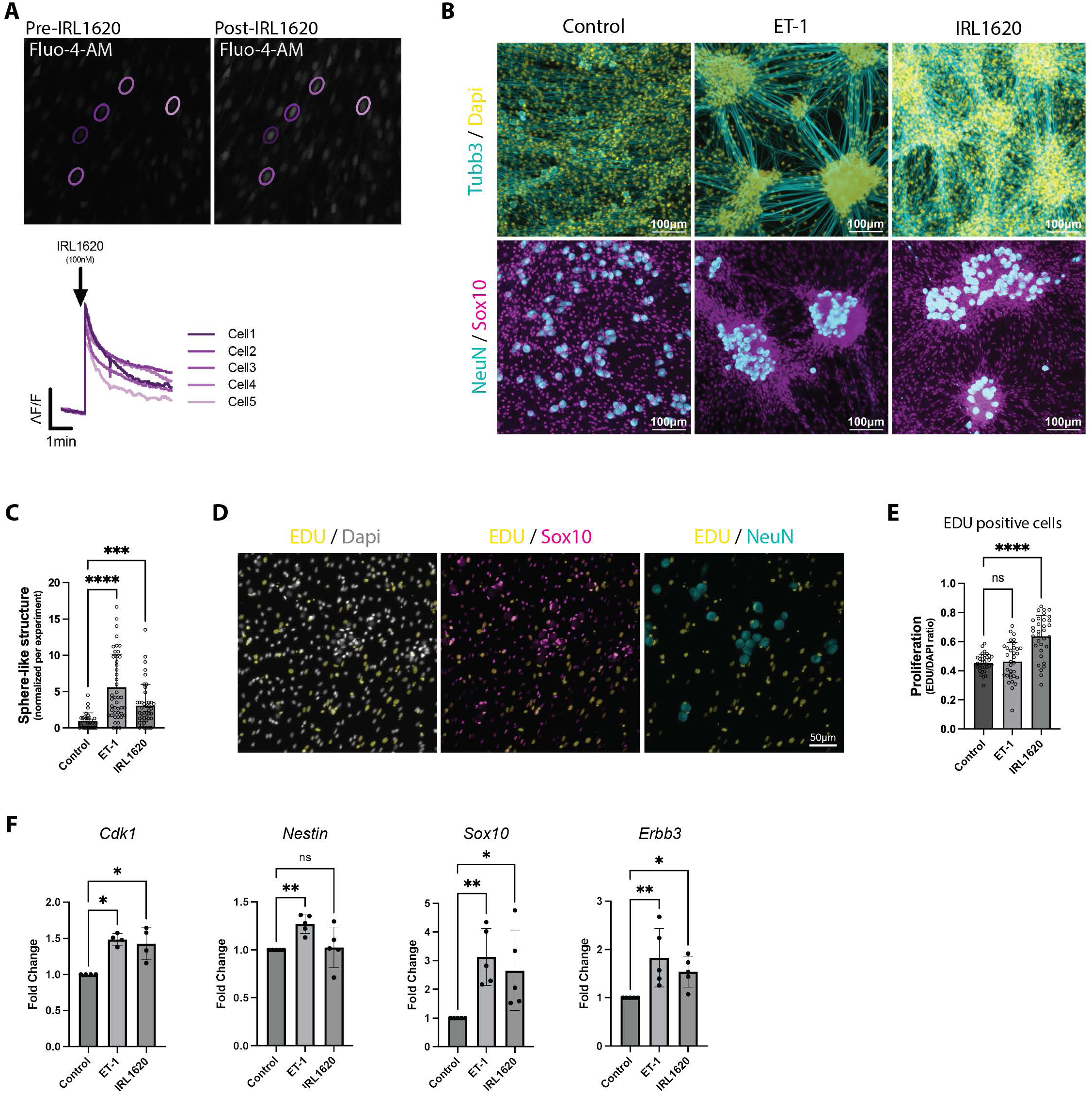
Activation of ETBR regulates embryonic glial progenitor proliferation. (**A**) Images of Fluo-4-AM signaling in eDRG cultures and representative calcium imaging traces of cells in response to the EDNRB agonist IRL1620 (100 nM). Colors correspond to encircled regions of interest. (**B**) Representative immunofluorescence images showing (Top panel) Dapi (yellow), Tubb3 (Cyan), (Bottom panel) NeuN (cyan), and Sox10 (magenta) expression in eDRG culture after 5 days of treatment with ET-1, IRL1620 compared to control. Under treated conditions “spherical structures,” cellular aggregates of neurons and glial progenitors appear (minimum diameter of 100 µm). Scale bars: 100 µm. (**C**) Quantification of sphere-like structure in eDRGs culture treated with ET-1 or IRL1620 compared to control. n= 30–46 technical replicates from 5 biological replicates per condition. (**D**) Representative immunofluorescence images of control eDRG spot culture showing EdU (yellow), Sox10 (magenta), NeuN (cyan), and Dapi (gray) staining. EdU^+^ nuclei co-localize with So×10^+^ but not with NeuN cells. Scale bars: 50 µm. (**E**) Quantification of proliferation, EdU^+^ cell ratio in eDRGs culture treated with ET-1 or IRL1620 compared to control. n= 33-34 technical replicates from 4 biological replicates per condition. (**F**) eDRG culture RTqPCR analysis of *Cdk1, Nestin, Sox10, Erbb3* relative mRNA expression after ET-1 or IRL1620 treatment compared to control. n= 3 technical replicates from 5 biological replicates per condition. Stat *p<0.05, **p<0.01, ***p<0.001, ****p<0.0001 by one-way ANOVA (**C, E-F**).

To confirm the physiological relevance of these findings beyond embryonic development, we investigated the expression and function of *Ednrb* in the DRG of postnatal day one (P1) and adult mice. Although the architecture of the P1 DRG is less characterized than in adults, previous studies have suggested that neuronal soma are enveloped by SGCs shortly after birth ^50, 51, 52^. Consistent with these earlier findings, we observed Fabp7 and Pdpn positive SGCs closely ensheathing Tuj1 positive neuronal bodies (Figure 6A). With ISH, we confirmed that *Ednrb* is expressed in both P1 and adult DRGs, with *Ednrb* co-localizing with *Fabp7* around sensory neurons (Figure 6B)^42^. To evaluate the functional role of the endothelin signaling in these cells, we used primary SGC cultures from P1 and adult mice (Figure S17A,D) ^53^. These cultures were highly enriched for SGC-specific markers across age and treatment conditions (Figure S17B,E,G). Functionally, SGCs from both P1 and adult stages responded to the ETBR agonist IRL1620 with an increase in Ca2+ fluorescence (Figure 6C-D). We then tested whether ETBR activation triggers proliferation of SGCs. In P1 SGC cultures, acute treatment (24h) with either ET-1 or IRL1620 led to a significant upregulation of the proliferation gene markers *Cdk1* and *Mki67* (Figure 6E). This was further supported by EdU incorporation assays, which showed a significant increase of proliferating cells (Figure 6G). IF confirmed that these EdU+ nuclei co-localized with SGC markers Sox10 and Glutamine Synthetase (GS) (Figure S17C). In contrast, this proliferative effect was absent in adult SGCs. Neither acute (24h) or chronic (10 days) treatment with ET-1 or IRL1620 induced the expression of *Cdk1* or *Mki67*; in fact, chronic treatment led to a significant downregulation of these markers (Figure 6F). Consistent with these transcriptional data, no changes in proportion of EdU+ cells were observed in adult cultures under any treatment condition (Figure 6H), despite these cells being Sox10+ and GS+ (Figure S17F, H). Together, these results demonstrate that the ability of ETBR signaling to drive cell proliferation is restricted to the postnatal period in SGCs. This suggests an age-related loss of proliferative plasticity in adult SGCs. Alternatively, the ETBR pathway may undergo a functional shift from promoting division in early development to maintaining or reinforcing quiescence in adult SGCs.

**Figure 6.**
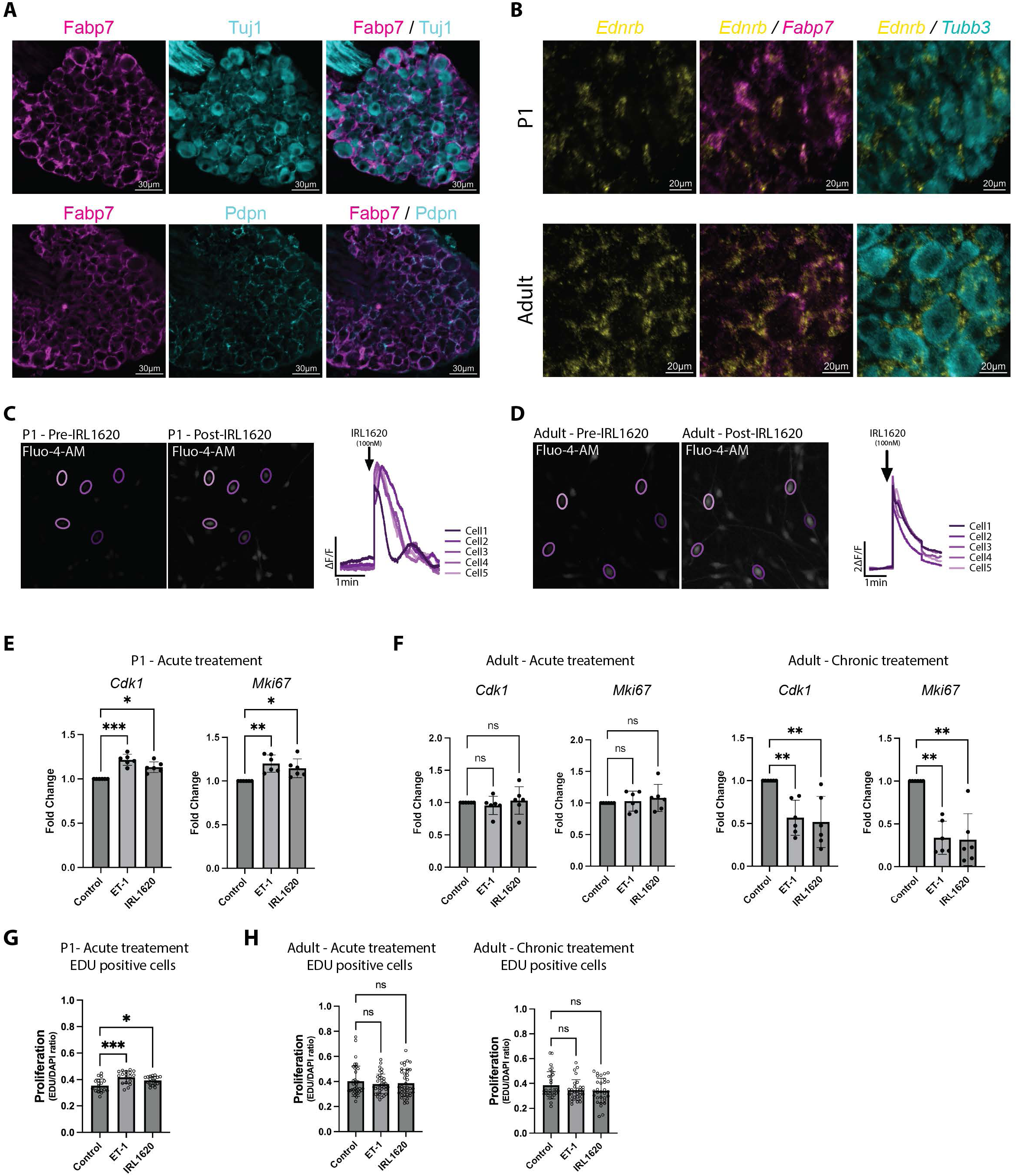
Endothelin signaling promotes glial proliferation during early postnatal stages. **(A)** Representative immunofluorescence images showing Fabp7 (magenta), Tuj1 (cyan - Top panel), Pdpn (cyan - Bottom panel). At P1, SGCs expressing Fabp7 and Pdpn surround neuronal soma. Scale bars: 30 µm. (**B**) Representative in situ hybridization images showing *Ednrb* (yellow) and *Fabp7* SGCs (magenta), and *Tubb3* neurons (cyan) on P1 (Top panel) and adult (Bottom panel) DRG section. Scale bars: 20 µm. (**C-D**) Images of Fluo-4-AM signaling in P1 (**C**) and adult (**D**) SGCs cultures and representative calcium imaging traces of cells in response to the EDNRB agonist IRL1620 (100 nM). Colors correspond to encircled regions of interest. (**E**) P1 SGCs culture RTqPCR analysis of *Cdk1, Mki67,* relative mRNA expression after 24h ET-1 or IRL1620 treatment compared to control. n=3 technical replicates from 6 biological replicates per condition. (**F**) Adult SGCs culture RTqPCR analysis of *Cdk1, Mki67,* relative mRNA expression after 24h (Left panel) or after 9 days (Right panel) ET-1 or IRL1620 treatment compared to control. n=3 technical replicates from 6 biological replicates per condition. (**G**) Quantification of proliferation, EdU^+^ cell ratio in P1 SGCs culture treated after 24h of ET-1 or IRL1620 treatment compared to control. n= 18 technical replicates from 4 biological replicates per condition. (**H**) Quantification of proliferation, EdU^+^ cell ratio in adult SGCs culture after 24h (Left panel) or after 9 days (Right panel) ET-1 or IRL1620 treatment compared to control. n= 30-38 technical replicates from 5 biological replicates per condition. Stat *p<0.05, **p<0.01, ***p<0.001 by one-way ANOVA (**E-H**).

## Discussion

In this study, we established a unified single-cell framework for peripheral glia by integrating diverse datasets across developmental stages and injury contexts. This approach resolves glial heterogeneity with high resolution and reveals a continuum of cellular states spanning homeostatic, reactive, and repair-associated programs. Within this landscape, we identified progenitor-like populations, localized within the Cajal’s IG, that occupy transitional positions within the glial lineage, with the potential to generate both myelinating and non-myelinating glial cells, and exhibit context-dependent expansion following injury. We also demonstrate that endothelin signaling via the ETBR receptor is a driver of glial proliferation during embryonic and early postnatal development, but not in adult stages. These findings provide a conceptual framework for understanding how glial plasticity is organized in the peripheral nervous system.

Primary sensory neurons exhibit profound diversity in morphology, physiology, and gene expression, which has long invited speculation that their surrounding SGCs may exhibit comparable heterogeneity. Our integrated atlas, comprising over 100,000 glial cell transcriptomes, provided an unprecedented opportunity to address this question. Surprisingly, high-resolution clustering failed to resolve SGCs into discrete transcriptional subtypes corresponding to neuronal classes. Instead, SGCs presented as a relatively uniform cell type at the transcriptional level, with only immune-responsive and IEG-expressing states distinguishable from the homeostatic population, consistent with prior studies ^1, 2, 21^. These IEG expressing SGCs may represent an active or poised state responding to neuronal signaling or a transient state induced by tissue dissociation. Despite the lack of distinguishing features at the transcriptional level, at the protein level, SGCs exhibit specialization correlating with the size of the neuronal subtype they envelop. The observed variability in marker expression at the protein level suggests that SGCs possess significant phenotypic plasticity, allowing them to adapt their molecular profiles to accommodate the unique metabolic and homeostatic demands of their associated neurons. In the CNS, astrocytes function as metabolic hubs that dynamically support neuronal activity ^54, 55^. They supply lactate, buffer extracellular calcium, clear glutamate to prevent excitotoxicity, and synthesize cholesterol for synaptic maintenance, acting as both bioenergetic suppliers and protective scaffolds ^54, 55^. Drawing a parallel to these CNS mechanisms, we propose that the heterogeneity observed in SGCs is not a result of fixed, intrinsic subtypes, but rather a reflection of a population adapting to the neuronal microenvironment. Notably, the enrichment of Glast (*Slc1a3*) and Kir4.1 (*Kcnj10*) in SGCs surrounding large-diameter neurons likely reflects an adaptation to the high-frequency firing patterns characteristic of these cells. Such heightened neuronal activity necessitates more efficient extracellular potassium buffering and accelerated glutamate clearance to maintain signal fidelity and prevent neurotoxicity. The higher abundance of Fabp7 in SGCs around large neurons also suggest that lipid metabolism pathways supply large neurons with lipids, such as cholesterol and phospholipids, to support neuronal function similarly to the role of astrocytes in the brain ^56, 57^. SGCs thus appear to function as a responsive metabolic hub that mirrors the specialized physiological requirements of their neuronal partners.

Another key finding of our study is the identification of a progenitor-like cell population characterized by expression of Oct6 (*Pou3f1*). Prior studies had identified such population ^1, 2, 3^, but low number of cells prevented detailed analyses and lacked examination of their localization within the DRG. We found that Oct6/*Pou3f1* cells are nested within the Cajal’s IG, consistent with recent studies in rat ^20^. In their studies, Koike et al 2025 proposed that SGCs in the distal part of the IG downregulate canonical markers (such as Fabp7, Kir4.1, and Kca2.3) and upregulate Oct6 (*Pou3f1*) to differentiate linearly along the axon into mSCs ^20^. Using high-resolution 3D imaging in mice, we found that SGCs create a continuous sheath around the neuronal cell body and proximal axon, supporting the presence of SGCs in the proximal Cajal’s IG in mice as well. However, our transcriptomic data and pseudotime trajectory analysis suggests a different model, in which *Pou3f1*+ cells represent a progenitor-like state. Rather than being restricted to a myelinating fate, these cells appear capable of generating either mSCs or SGCs/nmSCs. Together, these findings reframe Cajal’s IG as a niche harboring progenitors capable of adopting distinct fates depending on local cues. Identifying such cues will be of clinical relevance, since *Pou3f1/*Oct6 positive cells were also observed in human DRG ^20^.

Our analysis of injury conditions, incorporating the AIH paradigm alongside previously characterized models (DRC, SCI, SNC), revealed an unexpected dichotomy in glial transcriptional responses: cycling progenitors (G_CYC) only expanded following peripheral nerve injury, whereas post-mitotic progenitor-like cells (G_PROG) increased in abundance following central lesions. This selective engagement suggests that distinct injury contexts activate different progenitor pools within the DRG, potentially reflecting divergent regenerative demands. Notably, AIH, despite lacking direct physical trauma, resembled central injuries in both differential gene expression and cell abundance changes. This raises the intriguing possibility that hypoxic signaling engages transcriptional programs more closely aligned with central than peripheral injury responses, potentially through shared stress-response pathways independent of axonal damage.

Our results also identified ETBR signaling as a critical, though potentially temporally restricted driver of glial proliferation in the DRG. In embryonic and postnatal DRG cultures, ETBR activation triggered robust proliferation. However, this mitogenic capacity was apparently absent in adult SGCs, suggesting a significant age-related decline in glial plasticity. Several factors may account for this apparent loss of proliferative potential in the adult DRG. First, it is possible that only a discrete subpopulation of SGCs retains the capacity for ETBR-mediated proliferation in adult DRG, consistent with the limited renewal of SGC observed in adults ^17^. A small number of cells retaining proliferative responses to ETBR activation may fall below the sensitivity threshold of the methods we used in our culture model. Second, the absence of a proliferative response may be linked to cellular senescence. Indeed, ET-1 has been shown to trigger senescence in other cell types, such as myoblast, chondrocyte, endothelial cells and pericytes ^58, 59, 60, 61, 62^. Our previous findings of SGC atrophy in aged mice supports this hypothesis ^42^. Third, the downstream effectors of ETBR activation may be age and context dependent. ETBR is a pleiotropic receptor capable of coupling with various G-protein subunits ^63, 64^ potentially switching it signaling output from pro-proliferative to homeostatic or even senescence pathways in the adult. Such functional shifts could be further modulated by post-translational modifications, such as the metalloprotease-mediated cleavage of the ETBR N-terminal segment ^65, 66, 67^ or could be cell type and/or culture dependent condition ^68^. Fourth, cues in the environment may contribute to differential responses to ETBR activation. In the CNS, endothelin-1 (ET-1) signaling via ETBR regulate glial populations in the postnatal subventricular zone (SVZ), where it promotes the proliferation of radial glial cells and oligodendrocyte progenitor cells (OPC) while inhibiting their premature differentiation ^47^. However, in adult SVZ, ET-1 increases neural stem cell and OPC proliferation only following a demyelination injury ^47^. The proliferation of reactive astrocytes occurs following injury and requires the Notch1-STAT3-ETBR axis ^69^. Similar ETBR and Notch1 signaling mechanisms have been implicated in the survival and self-renewal of glioblastoma stem cells ^70, 71^. These findings suggest that the increase in progenitor abundance we observed following central lesions and AIH may result from increased ET-1 signaling and additional microenvironmental cues. Collectively, these findings suggest that the ETBR signaling axis is subject to complex temporal regulation, transitioning from a driver of development to a likely mediator of age-related glial remodeling.

In summary, this comprehensive atlas of individual cells establishes a molecular framework for the diversity and plasticity of peripheral glial cells. This framework resolves long-standing ambiguities in the classification of cell types within sensory ganglia. Identifying a population of progenitor-like cells in the specialized Cajal IG niche highlights a conserved structural domain for PNS plasticity. Additionally, our characterization of the endothelin signaling axis reveals a critical temporal shift in which ETBR-mediated proliferation may be restricted to the early developmental stage or injury conditions. This resource establishes a foundation for developing targeted strategies to promote repair and restore function in injured or diseased peripheral nervous systems.

## Data availability

The single-cell and single-nucleus RNA-sequencing data generated in this study have been deposited in the Gene Expression Omnibus (GEO) under accession GSE317728. The sciatic nerve crush time-course generated in this study have been deposited in GEO under accession GSE337372 (reviewer token: qvonqoiypvqzxwr). The Fmr1-KO DRG datasets was previously generated ^41^ and deposited under GSE176449. The processed integrated DRG and sciatic nerve cell atlas (Seurat object) is available at Zenodo (https://doi.org/10.5281/zenodo.21299570). Previously published datasets re-analyzed in this study are available under their original GEO accessions GSE236914, GSE139103, GSE158892, GSE198582, GSE213825, GSE142541, GSE201654, GSE175421, GSE182099, GSE198485, GSE147285, GSE137870 and GSE298413; the specific samples used are listed in Supplementary Table S1. Source data are provided with this paper.

## Code availability

All custom code used for atlas integration, clustering, annotation and downstream analyses (differential expression, gene co-expression network analysis, pseudotime, RNA velocity, reference mapping and label transfer, differential abundance, and integration assessment) is available at https://github.com/cavalli-lab-wustl/drg-sn-atlas).

## Acknowledgments

We would like to thank members of the Cavalli lab and Dr. Mayssa Mokalled for valuable discussions and suggestions. We gratefully acknowledge Dr. Peter Bayguinov for assistance with clearing and imaging tissues. Tissue clearing, imaging, and visualization were performed in part through the use of Washington University Center for Cellular Imaging (WUCCI) supported by Washington University School of Medicine, The Children’s Discovery Institute of Washington University and St. Louis Children’s Hospital (CDI-CORE-2015-505 and CDI-CORE-2019-813) and the Foundation for Barnes-Jewish Hospital (3770 and 4642). We would like to also thank Dr. Rui Feng and Dr. Eric E Ewan for their technical help with sample preparation for single cell data RNA sequencing. This work was funded in part by CRM Young Investigator Award to P.M, a Pilot Project Award from the Hope Center for Neurological Disorders at Washington University (V.C) and R21NS11549, R01 NS111719, R35 NS122260, DOD W81XWH-18-1-0729 to V.C. This publication is solely the responsibility of the authors and does not necessarily represent the official view of the National Institute of Health. The funders had no role in study design, data collection and analysis, decision to publish or preparation of the manuscript.

## Author Contributions

Conceptualization: PM, MT, VC; Data generation: PM, MT, OA, BT; Methodology: PM, MT; Sample processing: PM, OA, BT: Investigation: PM, MT; Visualization: PM, MT; Funding acquisition: VC, PM; Project administration: VC; Supervision: VC; Writing – original draft: PM, MT, VC; Writing – review & editing: PM, MT, OA, BT, VC

## Competing Interests Statement

The authors have no conflicts of interest or financial ties to disclose.

## Methods

### Experimental Animals & Surgical Procedures

All mice were approved by the Washington University School of Medicine Institutional Animal Care and Use Committee (IACUC) under protocol A3381-01. All experiments were performed in accordance with the relevant guidelines and regulations. All experimental protocols involving mice were approved by Washington University School of Medicine (protocol # 24-0078). Mice were housed and cared for in the Washington University School of Medicine animal care facility. This facility is accredited by the Association for Assessment & Accreditation of Laboratory Animal Care (AAALAC) and conforms to the PHS guidelines for Animal Care. Accreditation - 7/18/97, USDA Accreditation: Registration # 43-R-008.

All surgical procedures were performed under isoflurane anesthesia according to approved guidelines by the Washington University in St. Louis School of Medicine Institutional Animal Care and Use Committee. Adult female (8-12-week-old) C57Bl/6 mice were anesthetized using 2% inhaled isoflurane. Sciatic nerve injuries were performed as previously described ^25, 43, 72^. Briefly, the sciatic nerve was exposed with a small skin incision (∼1cm) and crushed for 5 seconds using #55 forceps. The wound was closed using wound clips and injured L4 and L5 dorsal root ganglia were dissected at the indicated time post-surgery. For spinal cord injury (SCI), a small midline skin incision (∼1cm) was made over the thoracic vertebrae at T9−T10, the paraspinal muscles freed, and the vertebral column stabilized with metal clamps placed under the T9/10 transverse processes. Dorsal laminectomy at T9/10 was performed with laminectomy forceps, the dura removed with fine forceps, and the dorsal column transversely cut using fine iridectomy scissors. Dorsal root injury was performed similarly as SCI, except that procedures were performed at the L2-L3 vertebral level, and fine forceps used to crush the right L4 and L5 dorsal roots for 5 seconds. L4 and L5 roots are in close proximity anatomically hence both roots were crushed simultaneously where the distance from the crush site to L4 DRG is 4-5mm and 7-8mm to L5 DRG. During dorsal root crush, the roots are forcefully squeezed causing the disruption of nerve fibers without interrupting the endoneurial tube. For AIH, we used a protocol that is within a low to moderate range ^73^, is sufficient to stabilize HIF-1α in DRG neurons and to induce expression of HIF-1α dependent genes ^74^ and is similar to an AIH protocol that was shown to induce motor plasticity in spared spinal pathways, improving respiratory and forelimb motor function in rats with chronic cervical injuries without inducing cell death ^75^. This protocol consists of alternating si× 10min episodes of hypoxia (8% oxygen) with si× 10min episodes of normoxia (21% oxygen) for a total of 120min (60min hypoxia), once daily for 3 days.

### Single-cell RNA sequencing

L4 and L5 DRG’s from mice 8-12 weeks old were collected into cold Hank’s balanced salt solution (HBSS) with 5% Hepes, then transferred to warm Papain solution and incubated for 20 min in 37°C. DRG’s were washed in HBSS and incubated with Collagenase for 20 min in 37°C. Ganglia were then mechanically dissociated to a single cell suspension by triturating in culture medium (Neurobasal medium), with Glutamax, PenStrep and B-27. Cells were washed in HBSS+Hepes +0.1%BSA solution, passed through a 70-micron cell strainer. Hoechst dye was added to distinguish live cells from debris and cells were FACS sorted using MoFlo HTS with Cyclone (Beckman Coulter, Indianapolis, IN). Sorted cells were washed in HBSS+, Hepes+ 0.1%BSA solution and manually counted using a hemocytometer. Solution was adjusted to a concentration of 500cell/microliter and loaded on the 10X Chromium system. Single-cell RNA-Seq libraries were prepared using GemCode Single-Cell 3′ Gel Bead and Library Kit (10x Genomics).

### Single-cell RNA sequencing data acquisition

Newly generated single-cell RNA sequencing libraries (2 acute intermittent hypoxia, 1 sciatic nerve crush 3 days post-injury, 1 naïve; GEO accession GSE317728) were prepared from adult mouse DRG as described above and sequenced on an Illumina NovaSeq 6000. Raw reads were aligned to the mm10 reference genome and count matrices were generated using Cell Ranger (v7.1.0, 10X Genomics). Previously published datasets were obtained from GEO as processed count matrices under accessions GSE236914, GSE139103, GSE158892, GSE198582, GSE213825, GSE142541, GSE201654, GSE175421, GSE182099, GSE198485, GSE147285, and GSE137870. Sample selection criteria, including specific conditions used from each accession, are detailed in Table S1.

### Data integration and batch correction

All analyses were performed in R (v4.3.3) using Seurat (v5.1.0). Count matrices were loaded and merged into a combined Seurat object and cells with >10% mitochondrial gene content were removed. Data were normalized using SCTransform, followed by PCA (50 components). Batch correction was performed using reciprocal PCA (RPCA) integration via IntegrateLayers with default parameters. A shared nearest neighbor graph was constructed using 50 integrated dimensions, and Louvain clustering was performed. UMAP was used for visualization.

### Iterative clustering and cell type annotation

Initial clustering identified seven major cell classes based on established marker genes: neurons (*Snap25, Syt1*), glial cells (*Sox10, Plp1*), fibroblasts (*Col6a1, Col1a1*), endothelial cells (*Pecam1, Flt1*), immune cells (*Aif1, Ptprc*), mural cells (*Rgs5, Acta2*), and erythrocytes (*Hba-a1*). Clusters with fewer than 1,000 mean detected genes (excluding erythrocytes) or poor representation across studies (<5 cells from all but one study) were removed. Each cell class was subset and re-integrated independently using the same RPCA pipeline. Clustering resolution was optimized per class using clustree, and clusters were annotated at major and minor cell type levels based on differential marker gene expression. Clusters with off-target class marker expression (multiplets), fewer than 1,000 mean detected features, or (for neurons) derivation from a single study (>95% of cells) were excluded. This hierarchical approach identified 7 cell classes, 28 major cell types, and 84 minor cell types.

### Differential gene expression analysis

Differential expression analysis was performed using FindMarkers in Seurat (Wilcoxon rank-sum test). Genes detected in ≥10% of cells in either group with log2 fold-change ≥0.5 (class-level) or ≥0.25 (within-class) were tested. P-values were adjusted using the Benjamini-Hochberg method (FDR < 0.05).

### Weighted gene co-expression network analysis

WGCNA was performed on glial cells using hdWGCNA (v0.3.03). Genes expressed in ≥3% of cells were retained. Metacells were constructed by aggregating neighboring cells (k = 25, maximum 10 shared cells) grouped by cluster identity within the integrated PCA space. A signed co-expression network was constructed with soft power automatically selected. Module eigengenes were harmonized across studies using Harmony. Hub genes were defined by eigengene-based connectivity (kME), and gene scoring used the top 30 hub genes per module.

### Gene Ontology enrichment analysis

GO enrichment analysis was performed using enrichGO from clusterProfiler (v4.10.1) with org.Mm.eg.db. Analysis was restricted to Biological Process terms. P-values were adjusted using the Benjamini-Hochberg method (FDR < 0.05).

### Pseudotime trajectory analysis

Trajectory analysis was performed on the G_PROG population using Slingshot (v2.10.0) with the integrated PCA embedding and default parameters. Spearman correlations between gene expression and pseudotime were calculated, and p-values were adjusted using the Benjamini-Hochberg method (FDR < 0.05). Expression dynamics were visualized using loess smoothing.

### Monocle3 pseudotime analysis

As an orthogonal trajectory method, the glia object was converted to a cell_data_set retaining the integrated UMAP, and a principal graph was learned with Monocle3 (version 1.4.26). Cells were ordered from a root defined as the G_PROG cell with maximal *Pou3f1* expression. To avoid graph over-branching at full scale, the analysis was restricted to the four major glial identities (G_PROG, SGC, mSC, nmSC) and balanced down-sampled to 4,000 cells per cluster (3,000 for the G_PROG-only graph). Genes varying along the graph were identified with graph_test (q < 0.05), and their per-gene pseudotime correlations were compared with the Slingshot ordering (Spearman ρ).

### RNA velocity analysis

RNA velocity analysis was restricted to Cavalli lab samples only. Spliced and unspliced transcript counts were quantified per library and combined with the glia object. RNA velocity was estimated with velocyto.R (v0.6). Per-cell velocity magnitude was computed as the L2 norm of the inferred expression change and compared across the naïve and injury conditions (SNC, DRC, SCI, AIH). Velocity-driver genes were ranked by mean absolute change and summarized by Gene Ontology enrichment (clusterProfiler).

### Reference mapping and label transfer

To demonstrate the utility of the atlas as a community reference, we mapped independent query datasets that were not used in atlas construction. Query cells were log-normalized and transfer anchors were identified against the integrated reference using FindTransferAnchors. Atlas Class, Major, and glia Minor identities were transferred, and query cells were projected into the reference UMAP using MapQuery, yielding per-cell predicted identities and prediction scores. To emulate a typical end-user workflow, query cells were additionally re-clustered on their own coordinates (SCTransform, RunPCA with 30 components, Harmony batch correction across samples, FindClusters resolution 0.5) and overlaid with atlas-predicted labels. Reference–query marker concordance was assessed by correlating the mean expression (AverageExpression, log1p) of the top 30 markers per cluster, reporting Pearson and Spearman coefficients. Query datasets comprised an Fmr1-KO DRG dataset (wild-type and knockout ^41^; GEO GSE176449), a sciatic nerve crush time-course (1, 7, 14, and 28 days post-injury, two replicates each; GEO GSE337372), and an aged DRG dataset (adult and aged; PIP-seq; ^42^; GEO GSE298413).

### Differential abundance analysis

Differences in cell-type composition were tested with propeller (speckle package, version 1.2.0) on arcsine-square-root–transformed cluster proportions (transform = “asin”), using the sample of origin as the replicate unit and testing at the Major and glia Minor levels. Two-group comparisons (for example, Fmr1-KO versus wild-type) used a moderated t-test; multi-group comparisons (injury time-course, aging) used an F-test followed by pairwise two-group tests. For atlas-wide injury-versus-control comparisons, robust empirical Bayes estimation was enabled.

### Integration assessment

Integration performance was quantified with the Local Inverse Simpson’s Index (LISI, version 1.0) computed on the full atlas without subsampling. Batch mixing (iLISI; higher values indicate better mixing) was scored across study, sequencing technology, tissue, and preparation (single cell versus single nucleus), and cell-type purity (cLISI; lower values indicate better preservation) was scored across Major cell types, comparing the unintegrated and integrated UMAP embeddings.

### Embryonic DRG cultures

DRG were isolated from time pregnant E13.5 CD-1 mice into dissection media consisting of DMEM (Gibco, Catalog #11-965-092) and Pen/Strep (ThermoFisher, Catalog #15140122). After a short centrifugation, dissection media was aspirated, and cells were digested in 0.05% Trypsin-EDTA (ThermoFisher, Catalog #25300054) for 25 min. Next, cells were pelleted by centrifuging for 2 min at 500 × g, the supernatant was aspirated, and Neurobasal was added. Cells were then triturated 25x and added to the growth medium containing Neurobasal (ThermoFisher, Catalog #21103049), B27 Plus (ThermoFisher, Catalog #A3582801), 100 ng/ml NGF (Alomone, Catalog #N-100), L-Glutamine (Gibco, Catalog #25030081), and Pen/Strep. Approximately 7,000 cells were added to each well in a 2.5 μl spot. Spotted cells were allowed to adhere for 10 min before the addition of the growth medium. Plates were pre-coated with poly-D-lysine and laminin (Poly-D-Lysine: Gibco, Catalog #A38904-01; Laminin: MilliporeSigma, Catalog #L2020.)

For drug treatment, ET-1 (Sigma-Aldrich, Catalog #E7764) 20nM, IRL1620 (Sigma-Aldrich, Catalog #SCP0135) 100 nM or vehicle control were added directly to the complete medium at DIV1 and incubated until DIV6.

For calcium assays, eDRG cells were plated on glass bottom 48 wells plates. Cells were washed twice and incubated in Neurobasal without red phenol (ThermoFisher, Catalog #12348017) for 20 min at 37C. Next, cells were incubated in 2 μM Fluo4-AM (ThermoFisher, Catalog #F14201) for 30 min, washed twice and placed in Neurobasal without red phenol for 10min at 37C. While at the microscope, cells were imaged for baseline fluorescence, followed by addition of 100nM IRL1620 (Sigma-Aldrich, Catalog #SCP0135) in Neurobasal without red phenol for time course imaging. Images were taken using an EVOS™ M7000. Analysis was performed in ImageJ (FIJI), cells analyzed and mean intensity of Fluo-4-AM fluorescence signal at baseline and over the time course was measured.

### SGCs cultures

We followed a previously established protocol for primary SGC cultures ^53^. Briefly, L1 and L5 DRGs were collected from P1 or 7-week-old mice and placed in cold dissection medium composed of HBSS (ThermoFisher, Catalog #14175–079) with 1% 1 M HEPES (ThermoFisher, Catalog #15630080). The DRGs were then transferred to freshly prepared pre-warmed dissociation medium containing 15 U/mL Papain suspension (Worthington Biochemical, Catalog #LS003126) 1% 1 M HEPES in HBSS. The samples were incubated at 37°C for 10 min for P1 samples and 20 min for adult samples. After washing the samples with pre-warmed HBSS, collagenase (150 µg/mL; Sigma, Catalog #C6885) was added, and the samples were incubated at 37°C for another 10min or 20 min for P1 samples or adult samples, respectively. Following wash with pre-warmed in complete media, the resulting single-cell suspension was gently triturated and resuspended in complete medium consisting of DMEM low glucose (ThermoFisher, Catalog #11885084) supplemented with FBS (Fisher scientific, Catalog #16140089), Pen/Strep (ThermoFisher, Catalog #15140122) and Amphotericin B (ThermoFisher, Catalog #15290026). The cell suspension was then passed through 50 µm and 20µm cell strainers successively. The single-cell suspension was then centrifuged at 300×rpm for 3 min, and the cell pellet was resuspended in complete medium. For P1 and adult acute drug treatment, ET-1 (Sigma-Aldrich, Catalog #E7764) 100nM, IRL1620 (Sigma-Aldrich, Catalog #SCP0135) 100 nM or vehicle control were added directly to the complete medium at DIV5 for P1 culture and DIV9 for adult culture for 24h. For adult chronic drug treatment, ET-1 100nM, IRL1620 100 nM or vehicle control were added directly to the complete medium at DIV1 and every 48h until DIV10.

For calcium assays, SGCs cells were plated on glass bottom 48 wells plates. Cells were washed twice and incubated in DMEM low glucose, without red phenol (ThermoFisher, Catalog #11054020) for 20 min at 37C. Next, cells were incubated in 2 μM Fluo4-AM (ThermoFisher, Catalog #F14201) for 30 min, washed twice and placed in DMEM without red phenol for 10min at 37C. While at the microscope, cells were imaged for baseline fluorescence, followed by addition of 100nM IRL1620 (Sigma-Aldrich, Catalog #SCP0135) in DMEM without red phenol for time course imaging. Images were taken using an EVOS™ M7000. Analysis was performed in ImageJ (FIJI), cells analyzed and mean intensity of Fluo4-AM fluorescence signal at baseline and over the time course was measured.

### Quantitative real-time PCR

For qRT-PCR, eDRG, and SGCs culture cells were collected, and total RNA was extracted using RNeasy Mini Kit (QIAGEN, Catalog #74104). For cDNA synthesis, 500 ng of RNA was converted into cDNA with the High-Capacity cDNA Reverse Transcription Kit (ThermoFisher, Catalog #4368814) according to manufacturer’s specifications. Quantitative PCR was completed using the PowerUp SYBR Green Master Mix (ThermoFisher, Catalog #A25780) using gene-specific primers (resource table) from Primerbank (https://pga.mgh.harvard.edu/primerbank/). qRT-PCR was performed on a QuantStudio 6 Flex System. Expression fold change for each gene of interest was calculated using the ΔCq method and normalized to the expression fold change of Gapdh expression compared to controls.

### Immunohistochemistry

For immunohistochemistry studies, mice were euthanized with CO_2_ asphyxiation, after isolation of mouse DRG, the tissue was postfixed using 4% paraformaldehyde for 24 hours at 4 degrees. PFA-fixed tissues were incubated in 30% sucrose in phosphate-buffered saline (PBS) overnight at 4°C, specimens were embedded in optimal cutting temperature compound (OCT) (Tissue-Tek), stored at −80°C until further processing. Frozen sections were cut at 10 µm mounted on Superfrost glass slides and stored at - 80°C. Sections were washed 3 times for 15 min each in 0.02 M PBS at room temperature. Sections were incubated overnight in 0.02 M PBS containing 0.3% Triton X100, and primary antibodies at room temperature in a humid chamber. The following day, sections were washed 3 times for 15 min each in 0.02 M PBS and incubated for 2h at room temperature with secondary antibodies, diluted in 0.02 M PBS containing 0.3% Triton X100. Then, sections were washed 3 times for 15 min in 0.02 M PBS and mounted using Vectashield medium containing DAPI (VectorLabs, Catalog #H-2000-10). Images were acquired using a Nikon TE2000E inverted microscope. Stained sections with only secondary antibody were used as controls. Fluorescence analysis was performed in ImageJ (FIJI).

For clearing, whole DRG samples were drop-fixed in 4% PFA for 30 minutes, post-fixed in 4% PFA at 4C° overnight, and washed in 0.02M PBS, 3 times for 15 min. Samples were then infiltrated with SHIELD polyepoxy ^76^ for 6 days with constant agitation at 4C°, were polymerized at 37C° for 24 hours, and were then delipidated using a SmartBatch+ ETC active tissue clearing platform (LifeCanvasTechnologies, Cambridge, MA). Samples were washed in 1x PBS for 24 hours and were permeabilized with 0.5% Triton X100 for 20min at room temperature. For staining, samples are incubated with primary antibodies diluted in 0.02M PBS, 0.3% Triton X100 for 8 hours at 37°C. After three 15-minute washes in 0.02M PBS, the sample are incubated with secondary antibodies diluted in 0.3% Triton X100 and 0.02M PBS for 3 hours at 37°C. After After three 15-minute washes in 0.02M PBS, samples were refractive index matched to 1.52 using EasyIndex RI matching media (LifeCanvasTechnologies). Whole sample images were acquired on a Nikon CSU W1 spinning disk microscope using a 10x/0.45 objective. Datasets were stitched together using NIS Elements (Nikon Instruments, Melville, NY) and were visualized using Bitplane Imaris 10.1.1 (Oxford Instruments, Abingdon, Oxfordshire).

For eDRG, SGCs cultures staining, cells were fixed in 4% PFA. Rise with PBS three times for 5 min, the fixed cells were blocked and permeabilized with 0.3% Triton X100 and blocked with 0.1% BSA in 0.02M PBS, 20 min at room temperature. For immunostaining following culture were incubated 3h in 0.02 M PBS containing 0.3% Triton X100, and primary antibodies at room temperature. The following day, sections were washed 3 times for 5 min each in 0.02 M PBS and incubated for 1h at room temperature with secondary antibodies, diluted in 0.02 M PBS containing 0.3% Triton X100 followed by incubation in 300 nM 4′,6-diamidino-2-phenylindole (DAPI; Sigma-Aldrich, Catalog #D9542) at room temperature for 10 min. Then, sections were washed 3 times for 5 min in 0.02 M PBS and kept in PBS until imaging. Cultured cells were imaged with acquired using the EVOS M7000 inverted microscope. Stained culture with only secondary antibody were used as controls. Cell counting analysis was performed in QuPath.

For the EdU experiment, cells cultures were treated with 10µM of EdU (Sigma-Aldrich, Catalog #BCK-EDU488, part No.CS219082) directly added to the complete media for 24h. First, the cells were fixed using 4% PFA diluted in PBS and incubated for 15 min at room temperature. After twice washes with 3% BSA in PBS, cells were permeabilized in 0.5% Triton X-100 in PBS, for 20 min at room temperature. After two washes with 3% BSA in PBS, cells were incubated in a fresh reaction cocktail (Sigma-Aldrich, Catalog #BCK-EDU488, part No.CS219085, CS219038, CS219083, CS219074) for 30 min at room temperature. Then, the cells were washed three times with 3% BSA in PBS before immunofluorescence staining. Cell counting analysis was performed in QuPath.

### RNAscope in situ hybridization

For RNAscope studies, mice were euthanized with CO2 asphyxiation, after isolation of mouse DRG, the tissue were incubated in 30% sucrose in phosphate-buffered saline (PBS) for 1h at 4°C, specimens were embedded in optimal cutting temperature compound (OCT) (Tissue-Tek), stored at −80°C until further processing. Frozen sections were cut at 14 µm mounted on Superfrost Plus glass slides and stored at - 80°C. The RNAscope fluorescent HiPlex V2 kit (Advanced Cell Diagnostics, ACD; Catalog #324419) was used according to the manufacturer’s instructions. Slides were retrieved from a –80°C freezer and immediately immersed in cold (4°C) 4% paraformaldehyde (PFA) for 60 min at room temperature. Tissues were dehydrated using a series of ethanol washes (50% ethanol for 5 min, 70% ethanol for 5 min, and 100% ethanol for 10 min) at room temperature. Slides were air-dried briefly, and hydrophobic boundaries were drawn around each section using a hydrophobic pen (ImmEdge PAP pen; Vector Labs). Once the boundaries were dry, protease IV reagent was added to each section and incubated for 30 min at room temperature. Slides were briefly washed in PBS at room temperature. Each slide was placed in a prewarmed humidity control tray (ACD) with dampened filter paper, and a mixture of probes was pipetted onto each section until fully submerged. The slides were then incubated in a HybEZ oven (ACD) at 40°C for 2 hr. Following the probe incubation, the slides were washed twice with 1 x RNAscope wash buffer and returned to the oven for 30 min after submersion in HiPlex Amp 1 reagent. The slides were washed twice with 1 x RNAscope wash buffer. This washing and amplification process was repeated using HiPlexAmp 2, and HiPlex Amp 3. Slides were then incubated with HiPlex Fluoro T1-T3 (Channel T1=Alexa 488, Channel T2= Dylight 550, Channel T3= Dylight 650). The slides were washed twice with 1x RNAscope wash buffer. Subsequently, the slides were processed with DAPI at room temperature for 30 seconds and cover-slipped using Prolong Gold Antifade mounting medium. DRG sections were imaged using a Nikon TE2000E inverted microscope.

## Supplemental information

**Figure S1.**
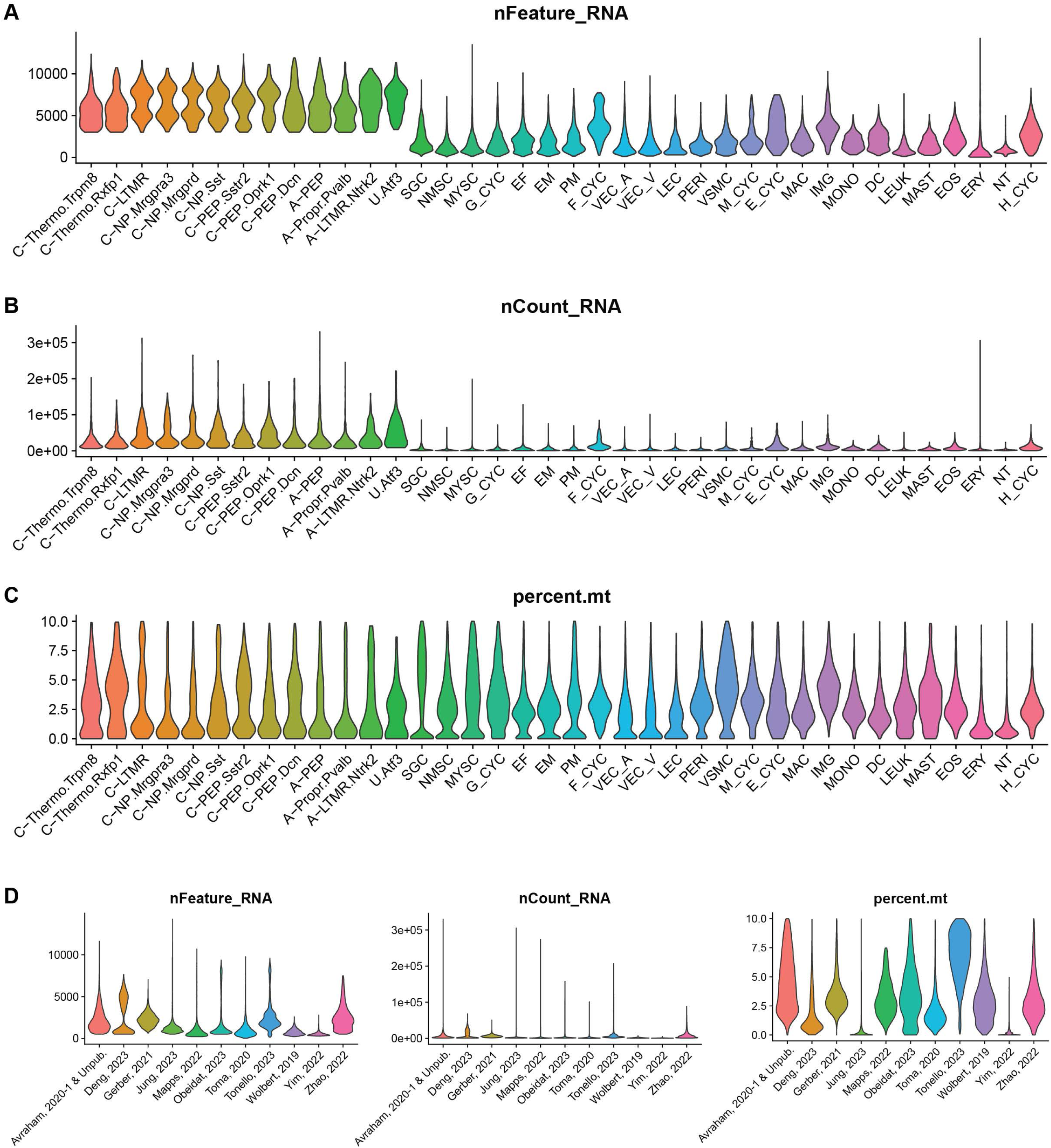
Quality control metrics for integrated dataset, related to Figure 1. (**A-C**) Violin plots showing the number of genes detected per cell (**A**), UMI counts per cell (**B**), and percentage of mitochondrial reads (**C**) across major cell types. Neuronal populations are shown at minor cell type resolution. (**D-F**) Violin plots showing the number of genes detected per cell (**D**), UMI counts per cell (**E**), and percentage of mitochondrial reads (**F**) across source studies.

**Figure S2.**
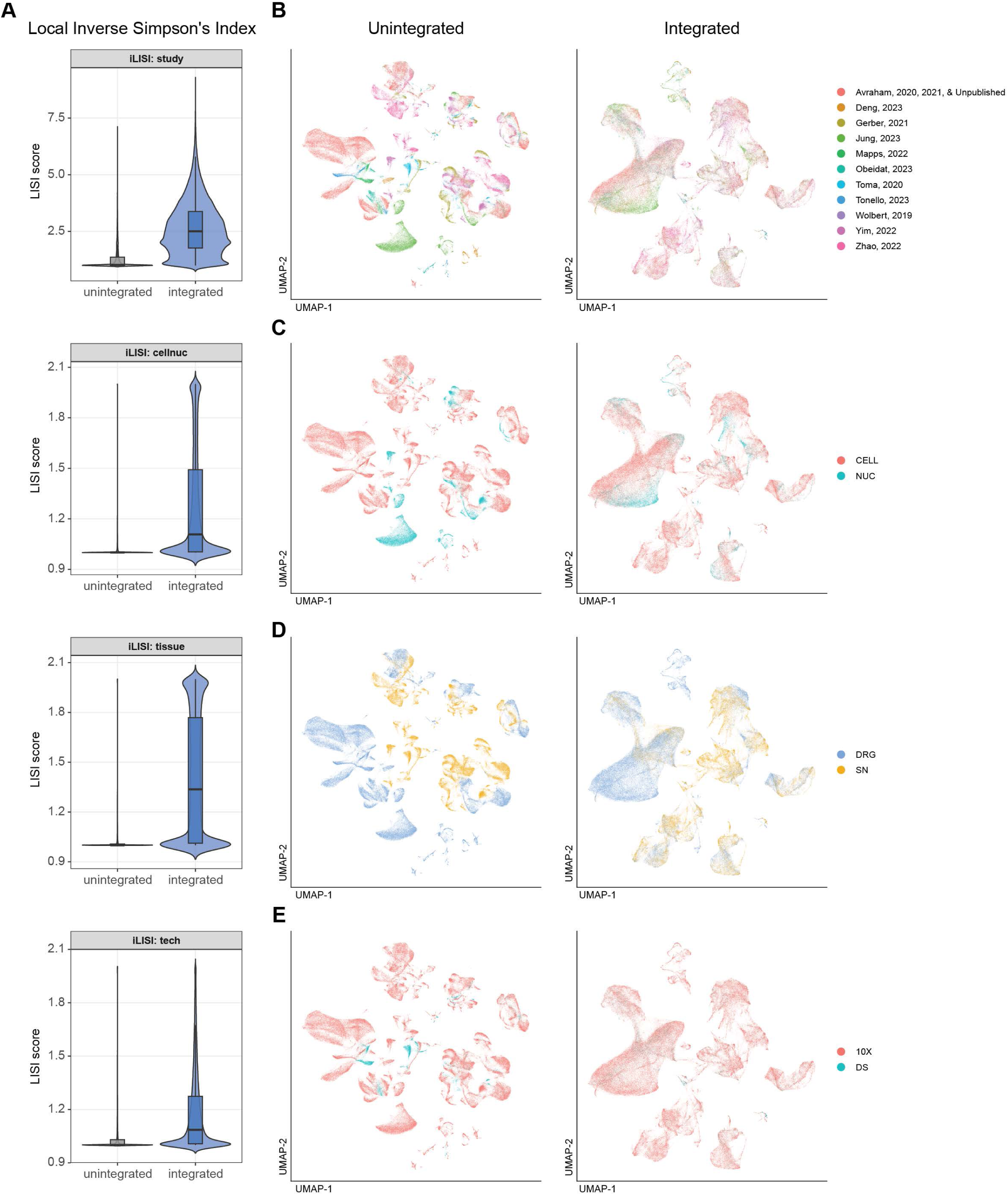
Integration efficacy assessment, related to Figure 1. (**A**) Local Inverse Simpson’s Index (LISI) score distributions quantifying mixing across study, sequencing preparation (cells vs. nuclei), tissue (DRG vs. SN), and technology, comparing pre-integration to post-integration. Higher LISI scores indicate better mixing across the indicated covariate. (**B-E**) Side-by-side UMAP visualizations of cells colored by study (**B**), sequencing preparation (**C**), tissue (**D**), and technology (**E**), shown for both the unintegrated (left) and integrated (right) embeddings, demonstrating that biological cluster structure is preserved while batch-driven separation is reduced.

**Figure S3.**
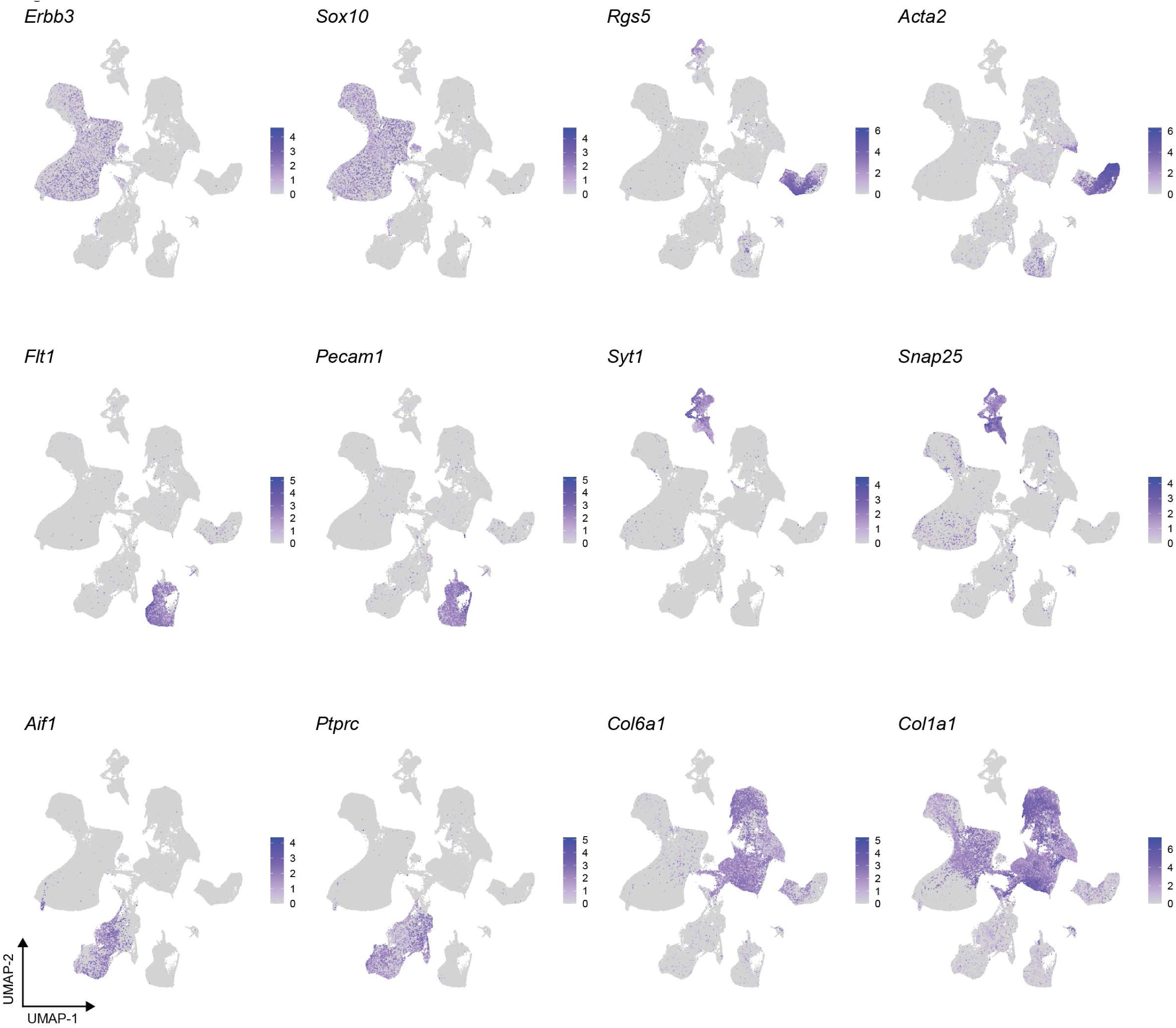
Expression of canonical cell type markers in integrated dataset, related to Figure 1. UMAP feature plots showing expression of marker genes used to annotate major cell types: *Erbb3* and *Sox10* (glia), *Rgs5* and *Acta2* (mural), *Flt1* and *Pecam1* (endothelial), *Syt1* and *Snap25* (neurons), *Aif1* and *Ptprc* (immune), *Col6a1* and *Col1a1* (fibroblasts).

**Figure S4.**
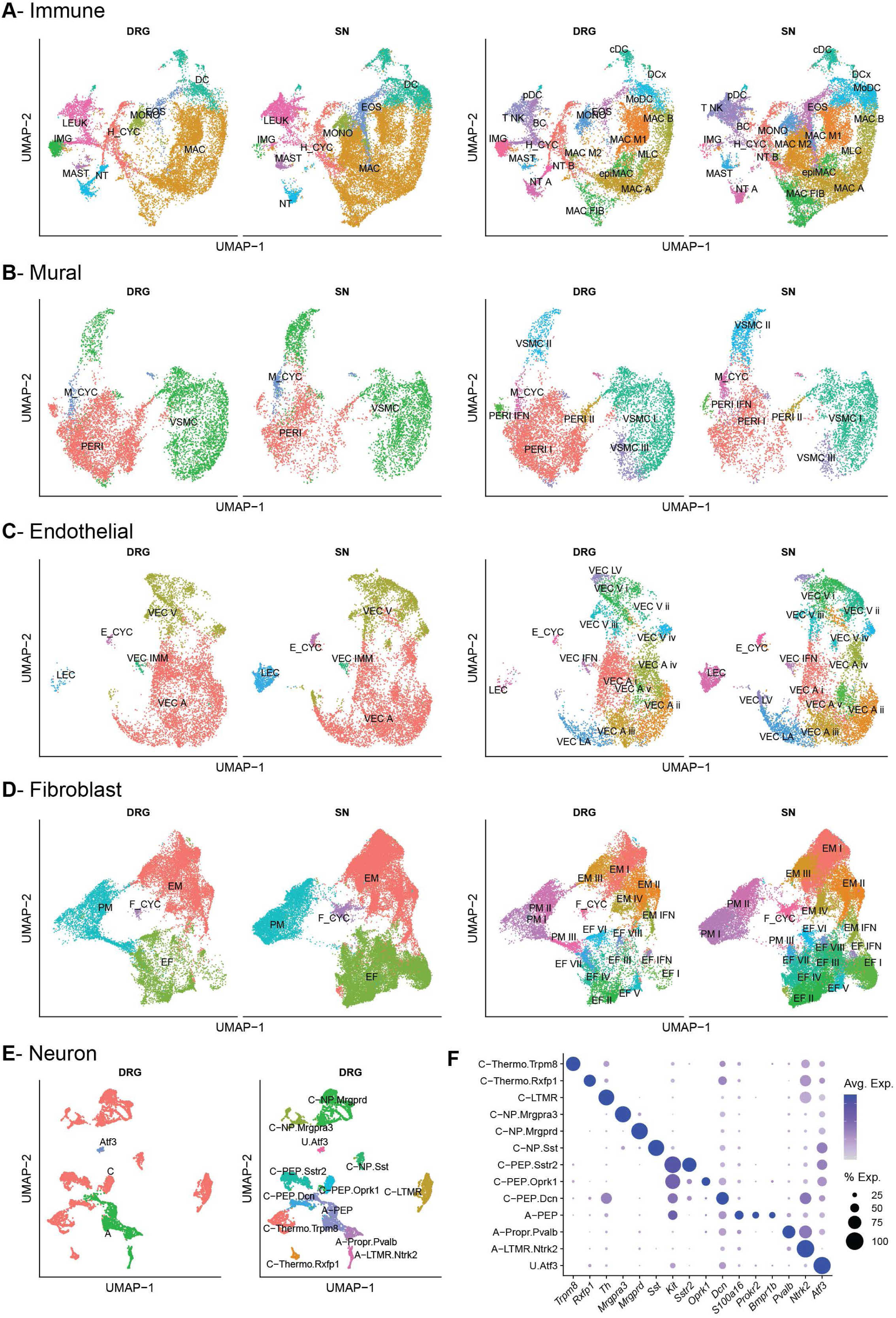
Subclustering and annotation of non-glial cell populations, related to Figure 1. (**A-D**) UMAP visualizations of cell class subsets: immune (**A**), mural (**B**), endothelial (**C**), and fibroblast (**D**) populations. For each class, cells are shown split by tissue (DRG, left; SN, right) and colored by major cell type (left pair) or minor cell type (right pair). Cell type abbreviations are defined in Supplemental File 1. (**E**) UMAP visualizations of DRG neuronal populations colored by major cell type (left) and minor cell type (right). Cell type abbreviations are defined in Supplemental File 1. (**F**) Dotplot of marker gene expression across minor neuronal subtypes. Dot size indicates percentage of cells expressing each gene; color intensity indicates average expression level.

**Figure S5.**
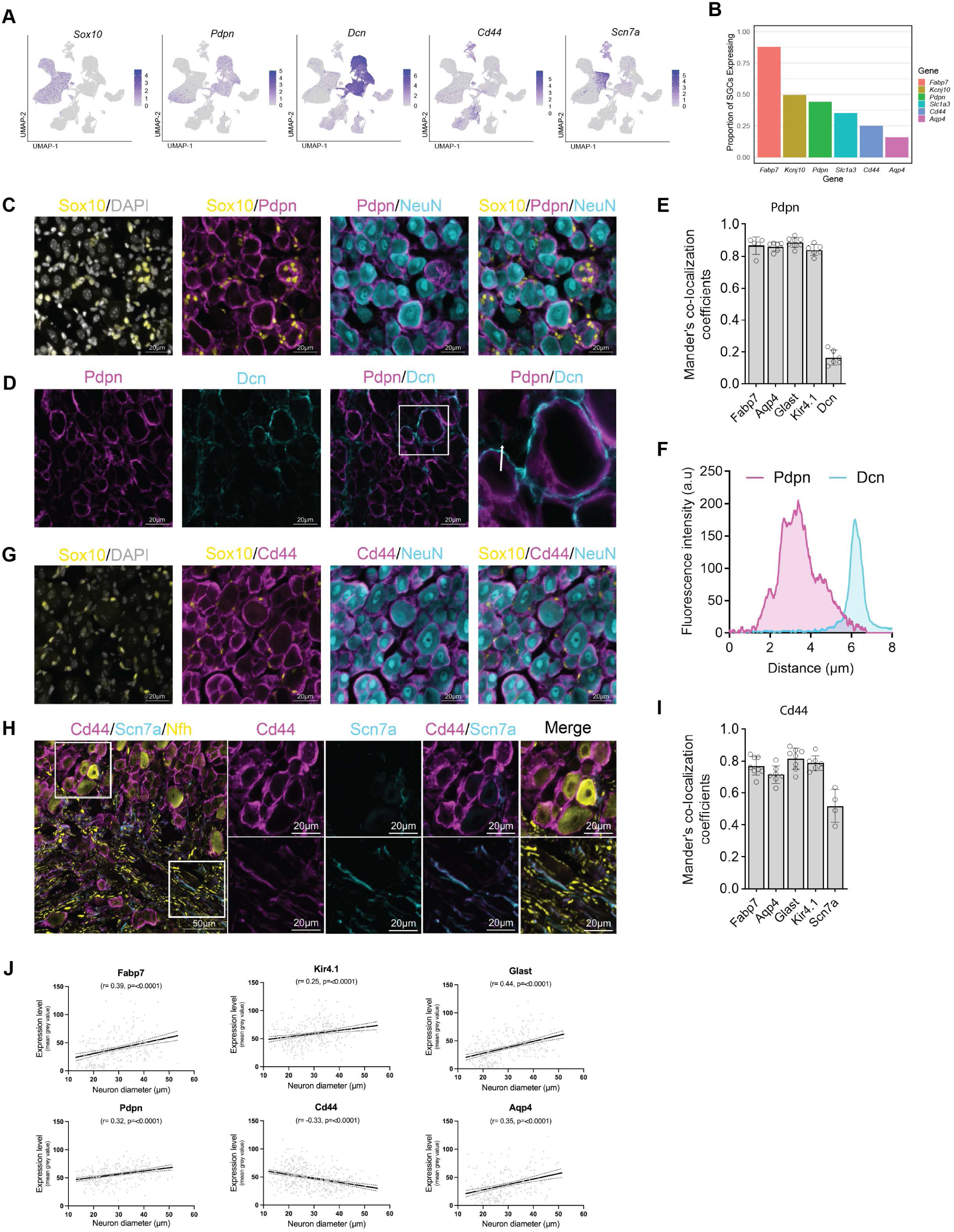
Molecular heterogeneity of SGCs and new marker characterization, related to Figure 2. (**A**) UMAP feature plots showing expression of genes *Sox10*, *Pdpn*, *Dcn* (fibroblast), *Cd44* and *Scn7a* (nmSC). (**B**) Bar plot showing the proportion of SGCs expressing *Fabp7, Kcnj10, Pdpn, Slc1a3, Cd44,* and *Aqp4*. (**C**) Representative immunofluorescence images showing Sox10 (yellow), Pdpn (magenta), NeuN (cyan), and DAPI (gray). Pdpn⁺ cells contain Sox10⁺ nuclei, surround NeuN⁺ neuronal soma. Scale bars: 20 µm. (**D**) Immunofluorescence labeling of Pdpn (magenta) and Dcn (cyan). Merged images and high magnification inset demonstrate fibroblast marker Dcn surrounding Pdpn⁺ SGCs. Scale bars: 20 µm. (**E**) Manders’ co-localization coefficients quantifying overlap between Pdpn and SGCs markers (Fabp7, Aqp4, Glast, and Kir4.1) or the fibroblast marker Dcn. Data are presented as mean ± s.e.m. Each dot represents one biological replicate. (**F**) Fluorescence intensity profiles measured along the arrow indicated in E, showing spatial segregation of Pdpn and Dcn expression. (**G**) Representative images showing Sox10 (yellow), Cd44 (magenta), NeuN (cyan), and DAPI (white). CD44⁺ cells contain Sox10⁺ nuclei, surround NeuN⁺ neuronal soma. Scale bars: 20 µm. (**H**) Low- and high-magnification immunofluorescence images showing Cd44 (magenta), Scn7a (cyan), and Nfh (yellow). Boxed regions indicate neuronal soma (top) Cd44⁺ SGC surround Nfh neuronal soma and axonal regions (bottom) Cd44 colocalizes with Scn7a⁺ nmSC. Scale bars: 50 µm (overview), 20 µm (high magnification). (**I**) Manders’ co-localization coefficients quantifying overlap between Cd44 and SGCs markers (Fabp7, Aqp4, Glast, and Kir4.1) and Scn7a. Data are presented as mean ± s.e.m. Each dot represents one biological replicate. (**J**) Scatter plots showing correlations between neuronal soma diameter and expression levels of Fabp7, Kir4.1 (*Kcnj10*), Pdpn, Glast (*Slc1a3*), Cd44, and Aqp4. Each dot represents an individual ring of SGCs surrounding a neuron; solid lines indicate linear regression fits with 95% confidence intervals. Pearson correlation coefficients (r) and p-values are shown.

**Figure S6.**
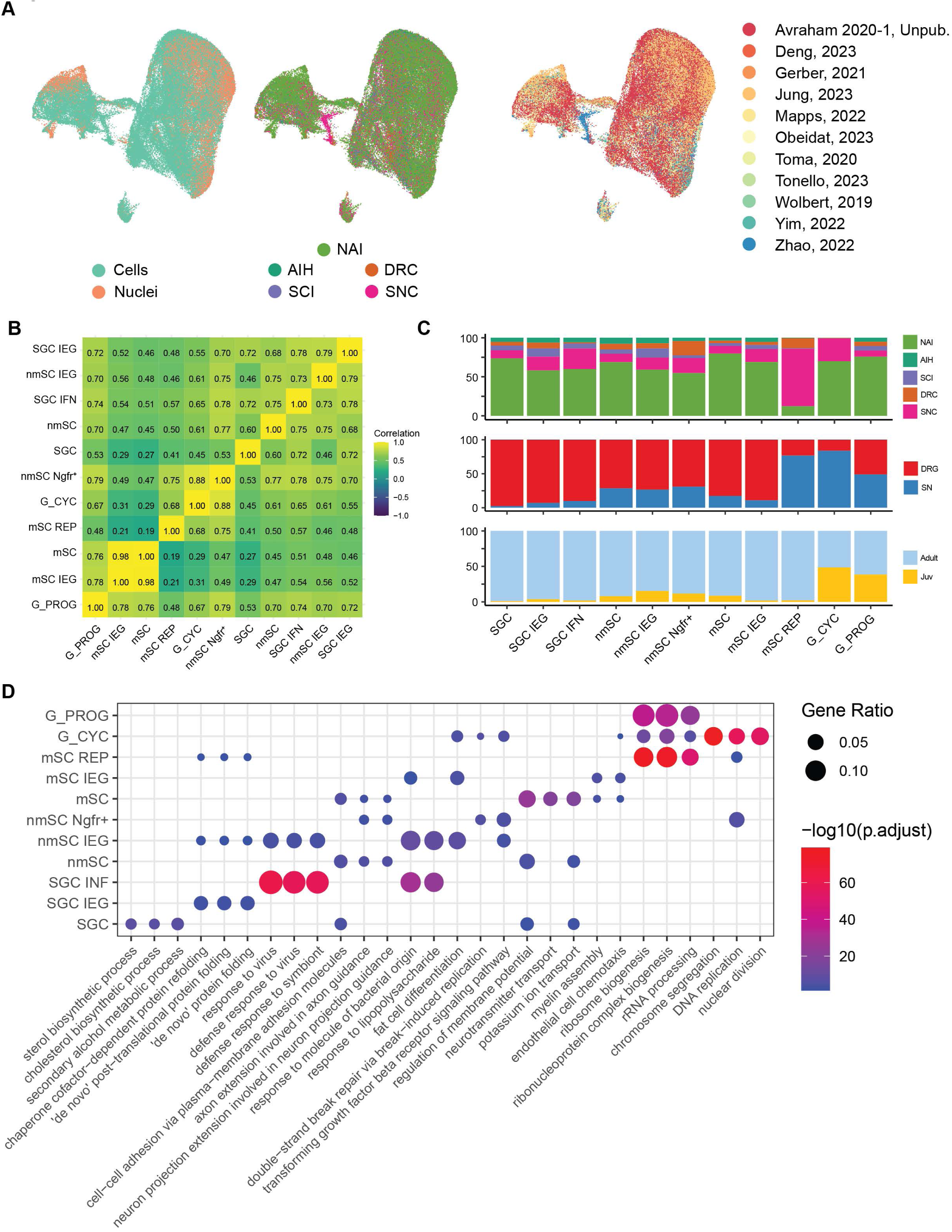
Characterization of glial subpopulation heterogeneity and functional enrichment, related to Figure 3. (**A**) UMAP visualizations of glial cells colored by sequencing preparation (single-cell or single-nucleus), treatment condition, and source study. (**B**) Correlation heatmap of glial subpopulations based on expression of the top 3,000 variable genes. (**C**) Stacked bar graphs depicting the relative proportion of cells within each glial subpopulation by treatment condition, tissue, and age. (**D**) Dotplot of Gene Ontology (GO) biological process terms enriched in each glial subpopulation based on cluster marker genes. Analysis was restricted to terms appearing in three or fewer populations. Dot size indicates gene ratio; color intensity indicates −log10(adjusted p-value).

**Figure S7.**
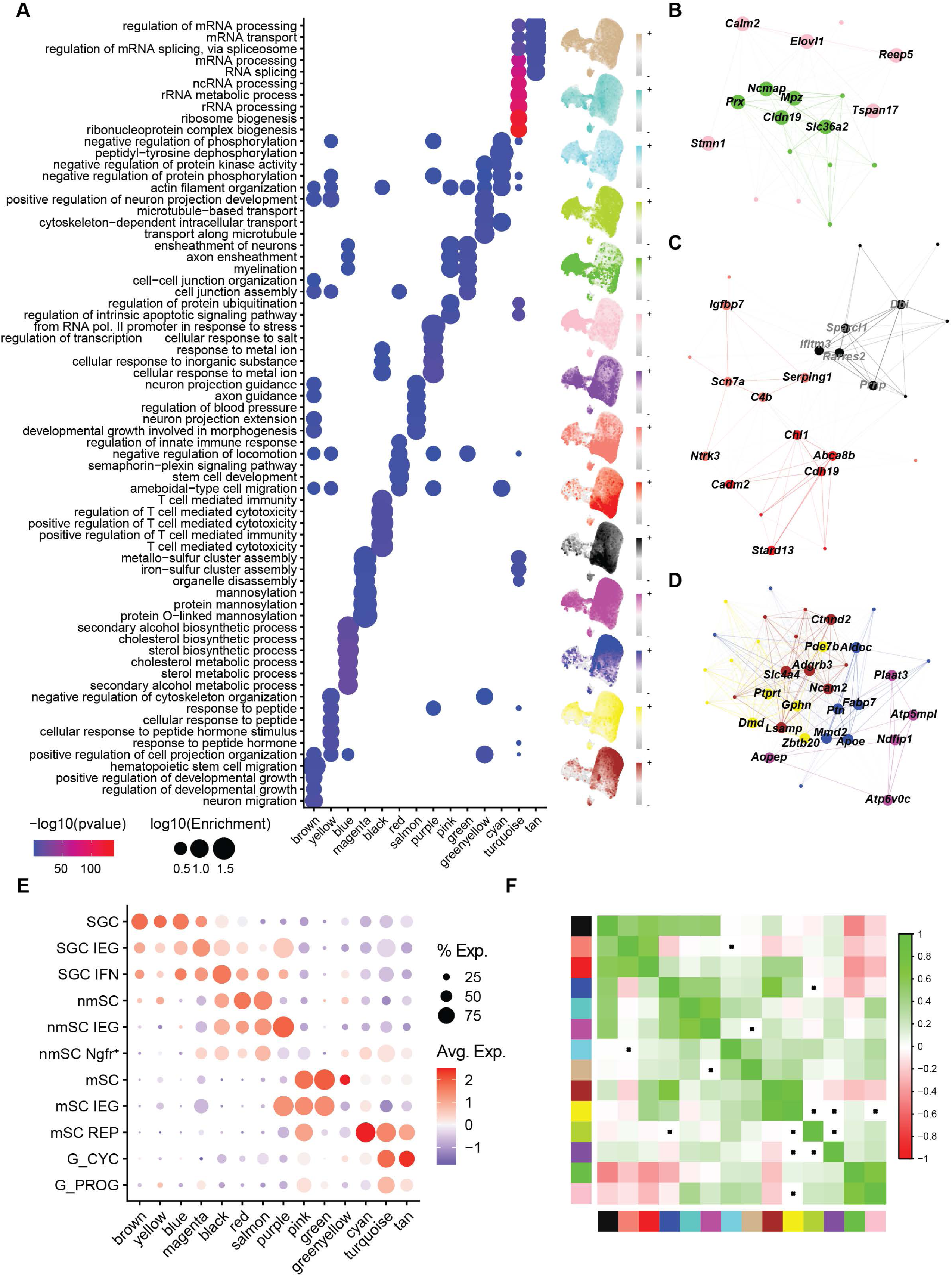
Weighted gene co-expression network analysis of peripheral glia, related to Figure 3. (**A**) Integrated visualization of Gene Ontology (GO) biological process terms enriched in each WGCNA module (left) with corresponding UMAP feature plots of module expression (right). Dot size indicates log10(enrichment); color intensity indicates −log10(p-value). (**B**) Network diagram of top hub genes in mSC-associated modules (green, pink). (**C**) Network diagram of top hub genes in nmSC-associated modules (black, red). (**D**) Network diagram of top hub genes in SGC-associated modules (yellow, brown, blue, magenta). (**E**) Dotplot of module expression across glial subpopulations. Dot size indicates percentage of cells expressing module genes; color intensity indicates average expression level. (**F**) Correlation heatmap of module eigengenes.

**Figure S8.**
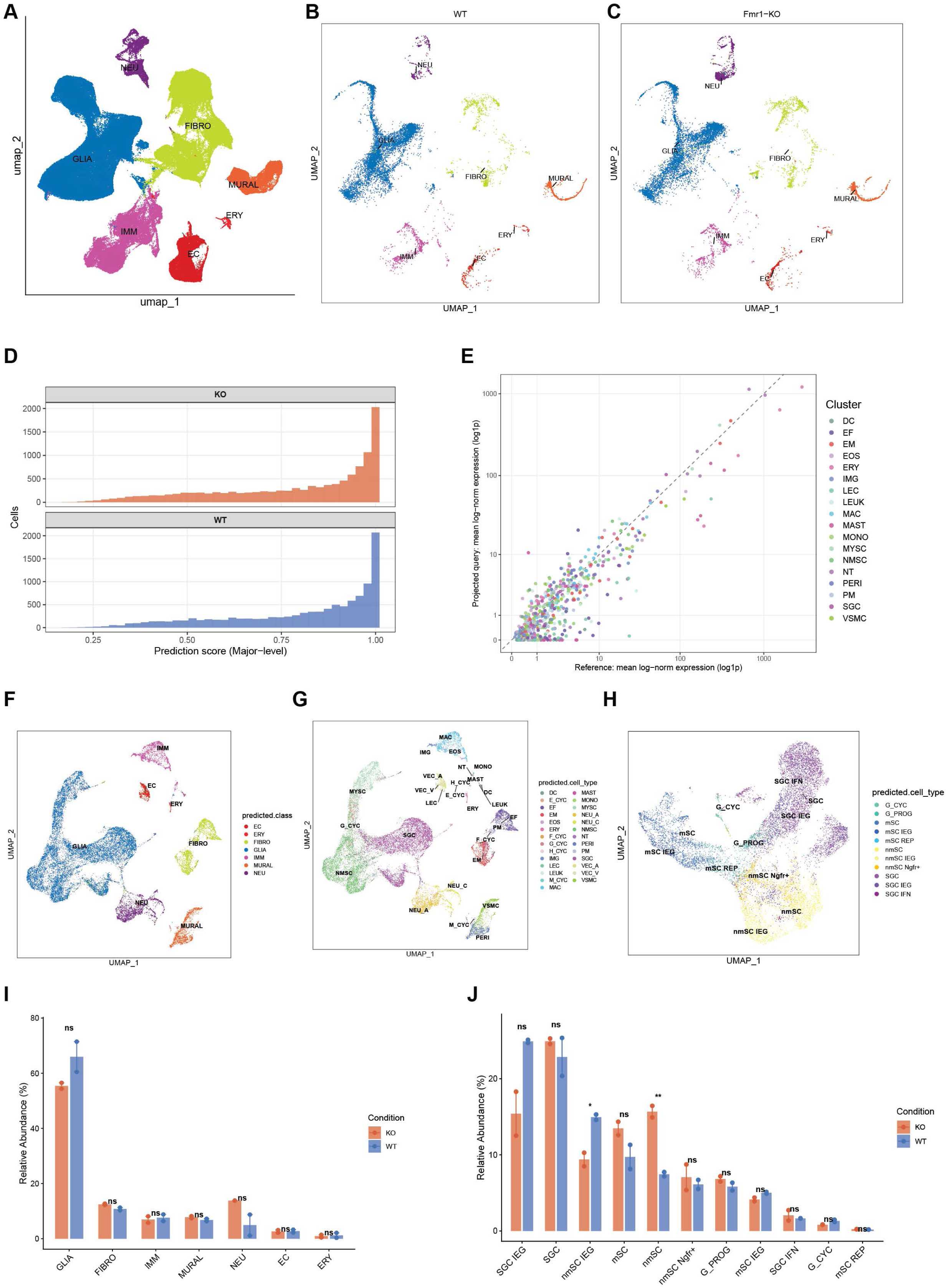
Demonstration of atlas utility — label transfer onto an independent Fmr1KO DRG dataset, related to Figure 1. (**A**) UMAP of the integrated reference atlas colored by the 28 Major cell types. (**B-C**) Query cells from wild-type (**B**) and Fmr1-KO (**C**) DRG projected onto the reference UMAP and colored by atlas-predicted Major cell type. (**D**) Major-level mapping confidence (prediction score) distributions for Fmr1-KO (top) and wild-type (bottom) cells. (E) Marker-expression concordance between reference and query for the top 30 markers per Major cluster (without label transfer); Spearman ρ = 0.78, Pearson R = 0.87. (**F-H**) Query cells re-clustered on their own coordinates (SCTransform + Harmony) and colored by atlas-predicted Class (**F**), atlas-predicted Major cell type (G), and, for the glia subset, atlas-predicted Minor cell type (**H**). (**I**) Class-level relative abundance comparison between Fmr1-KO and wild-type; statistical comparison by propeller on asin-transformed proportions (ns: not significant, * FDR<0.05, ** FDR<0.01, *** FDR<0.001). (**J**) Glia Minor-level relative abundance comparison; propeller statistics as in (**I**).

**Figure S9.**
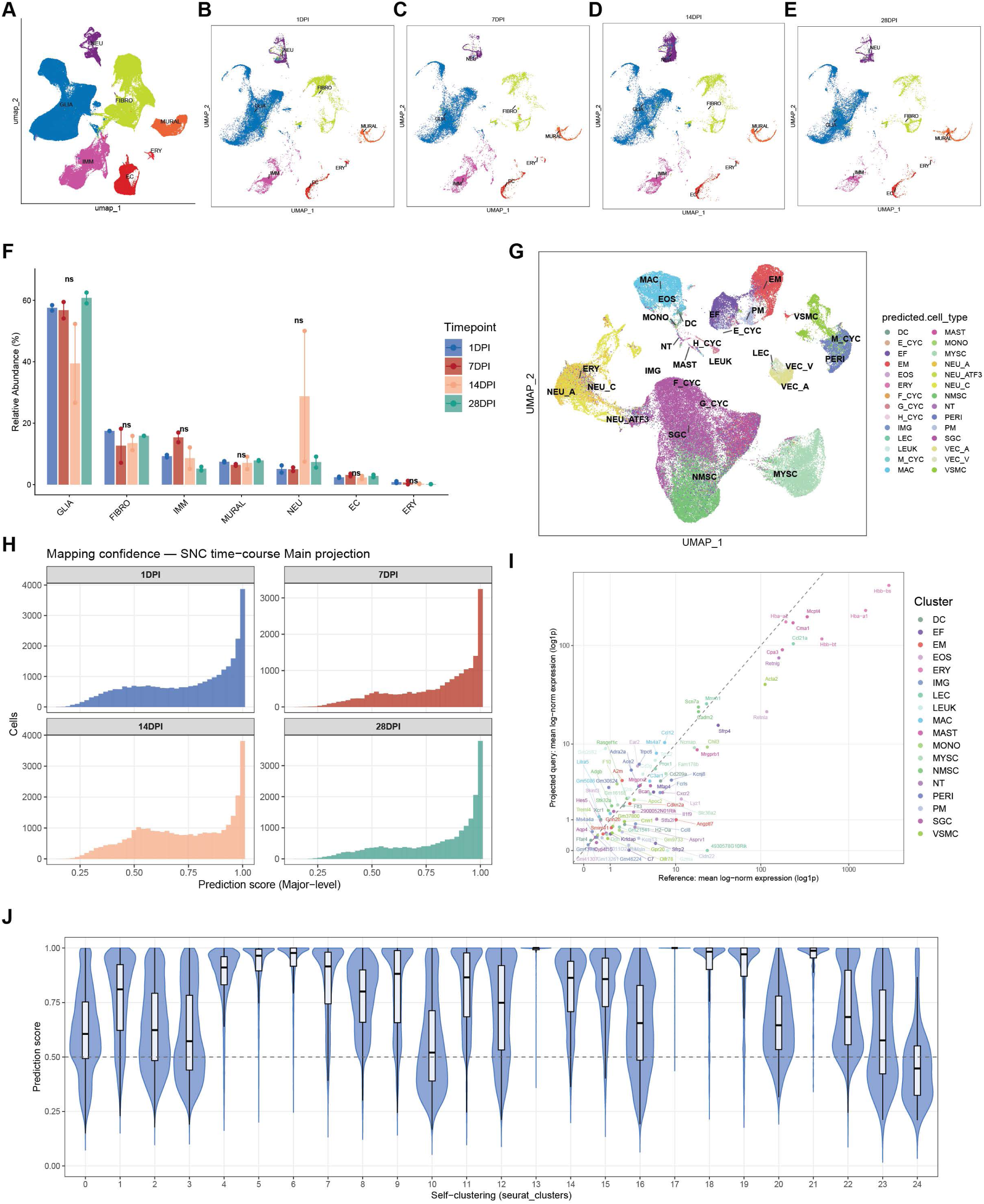
Demonstration of atlas utility — label transfer onto a sciatic nerve crush (SNC) time-course dataset, related to Figure 1. (**A**) Reference atlas UMAP. (**B-E**) UMAP projection of SNC time-course samples onto the reference atlas, split by post-injury day (1, 7, 14, 28 DPI), colored by atlas-derived Major cell type. (**F**) Major cell-type relative abundance across timepoints; statistical comparison by propeller F-test with asin-transformation. (**G**) Atlas-predicted Major cell-type assignments overlaid on the integrated UMAP. (**H**) Major-level mapping confidence histograms split by timepoint. (**I**) Marker concordance between atlas reference and projected labels at the Major level; Spearman ρ and Pearson R reported. (**J**) Atlas-label confidence per self-defined cluster (Major-level).

**Figure S10.**
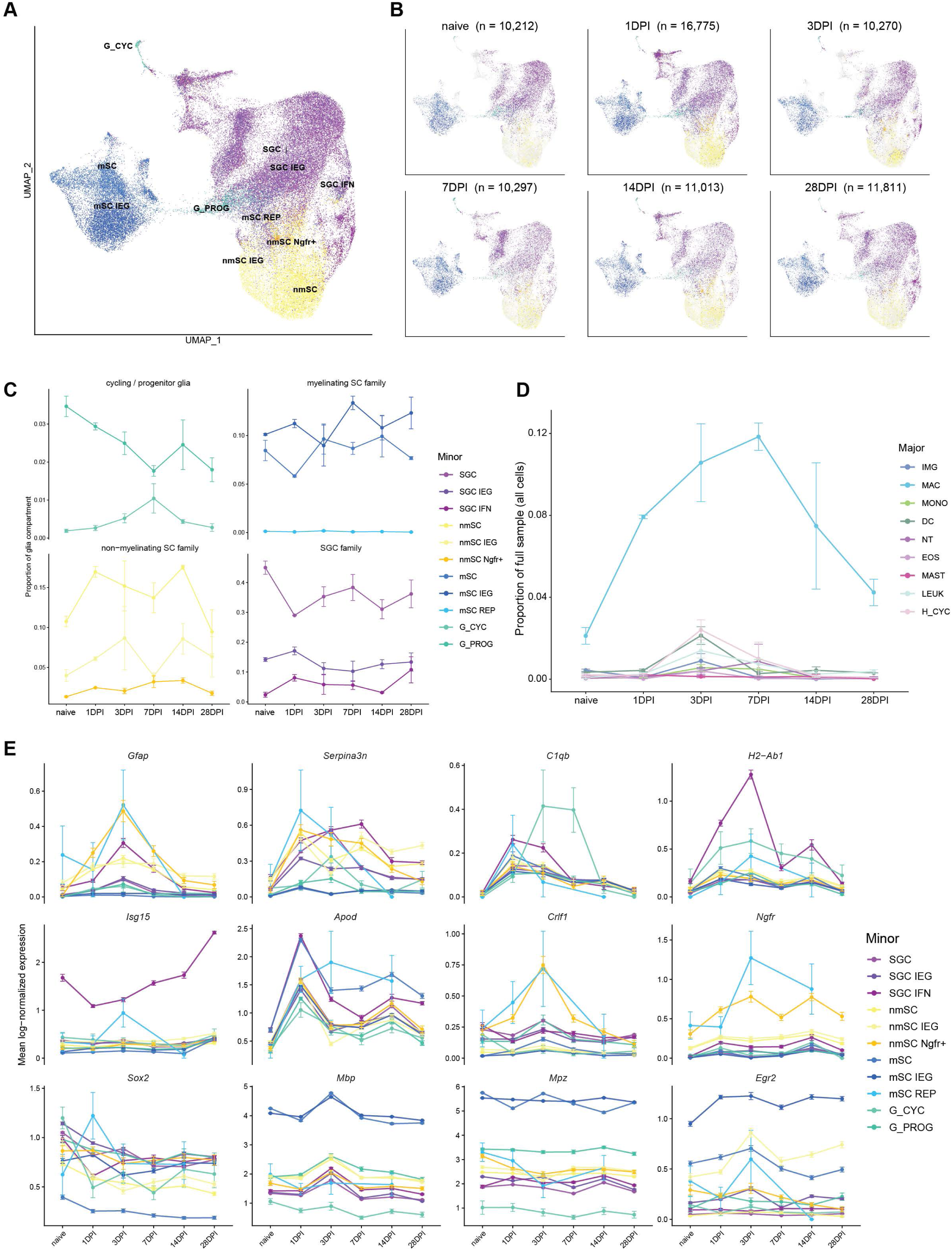
Sciatic nerve crush time-course — glial and immune cell dynamics, related to Figure 1. (**A**) UMAP of the glial compartment of the integrated sciatic nerve crush (SNC) time-course, colored by the 11 Minor glial cell types. (**B**) Per-timepoint density UMAPs of glial cells at naïve (n = 10,212), 1 (n = 16,775), 3 (n = 10,210), 7 (n = 10,297), 14 (n = 11,013), and 28 (n = 11,611) days post-injury (DPI). (**C**) Relative abundance trajectories across timepoints for four glial families — cycling/progenitor glia (G_CYC, G_PROG), myelinating Schwann cells (mSC, mSC IEG, mSC REP), non-myelinating Schwann cells (nmSC, nmSC IEG, nmSC Ngfr+), and satellite glia (SGC, SGC IEG, SGC IFN) — expressed as proportion of the glial compartment; points and error bars indicate mean ± SEM across replicate samples per timepoint. (**D**) Relative abundance trajectories of major immune cell populations (IMG, MAC, MONO, DC, NT, EOS, MAST, LEUK, H_CYC) across timepoints, expressed as proportion of all cells (mean ± SEM across replicate samples), highlighting the prominent macrophage (MAC) response that peaks within the first post-injury week. (**E**) Mean normalized expression trajectories across timepoints for representative glial-response genes (Gfap, Serpina3n, C1qb, H2-Ab1, Isg15, Apod, Crlf1, Ngfr, Sox2, Mbp, Mpz, Egr2), shown per Minor cell type; error bars indicate SEM across cells.

**Figure S11.**
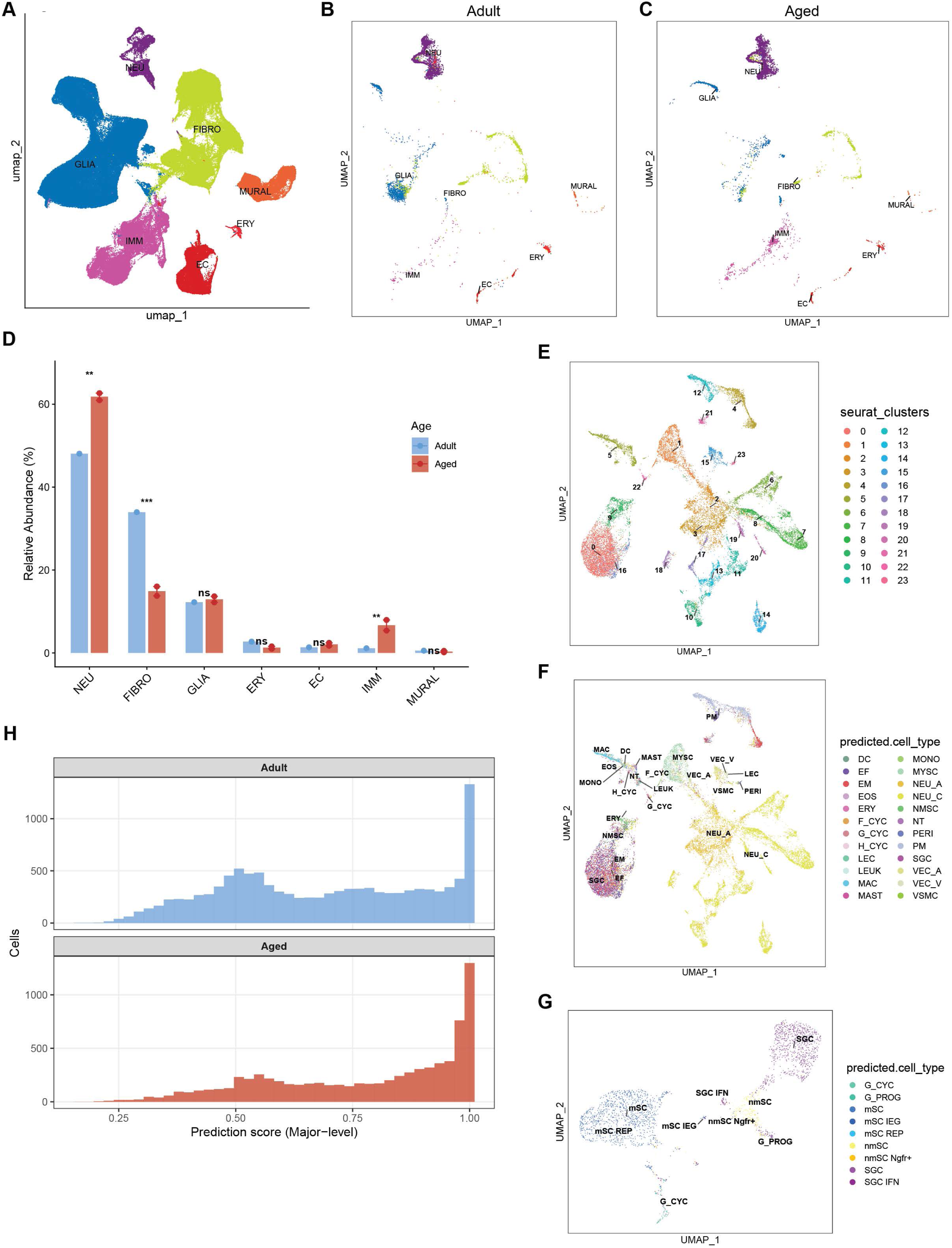
Demonstration of atlas utility — label transfer onto an aged DRG dataset (Feng et al., 2025; PIP-seq), related to Figure 1. (**A**) Reference atlas UMAP. (**B-C**) UMAP projection of Adult (**B**) and Aged (**C**) DRG cells onto the reference atlas, colored by atlas-derived Major cell type. (**D**) Major cell-type relative abundance comparison between Adult (n=1) and Aged (n=2) DRG samples; statistical comparison by propeller with asin-transformation. (**E**) Self-clustering UMAP of the aged dataset (Harmony integration, resolution 0.5) colored by self-defined cluster identity. (**F**) Atlas-predicted Major cell-type assignments overlaid on the self-clustering UMAP. (**G**) Atlas-predicted Minor cell-type assignments at the glia level. (**H**) Major-level mapping confidence histograms for Adult (top) and Aged (bottom) samples.

**Figure S12.**
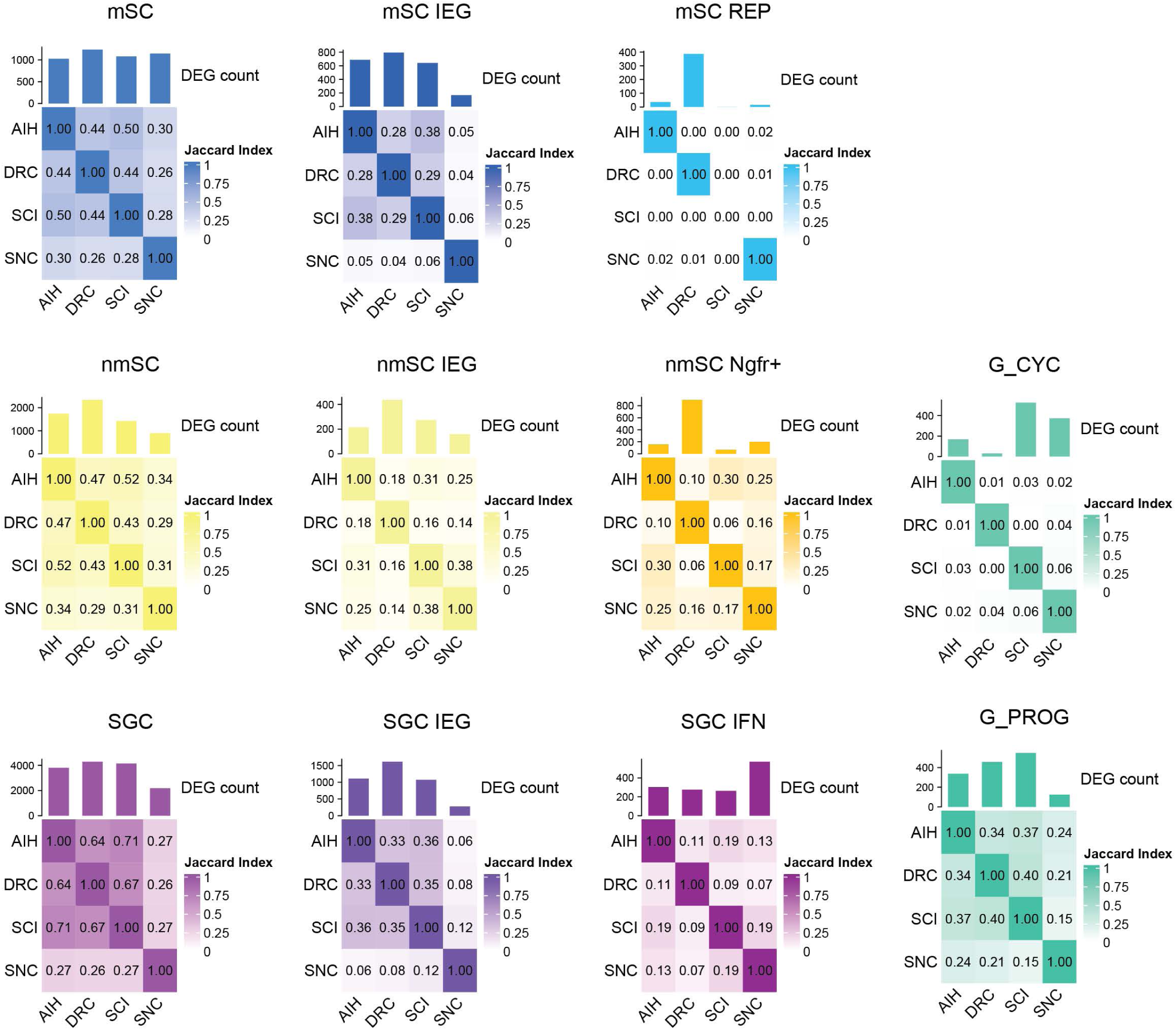
Transcriptional overlap of glial injury responses across injury paradigms, related to Figure 3. Jaccard similarity heatmaps showing pairwise overlap of differentially expressed genes between injury conditions (AIH, DRC, SCI, SNC) for each glial subpopulation. Bar graphs above each heatmap indicate total DEG count per injury condition.

**Figure S13.**
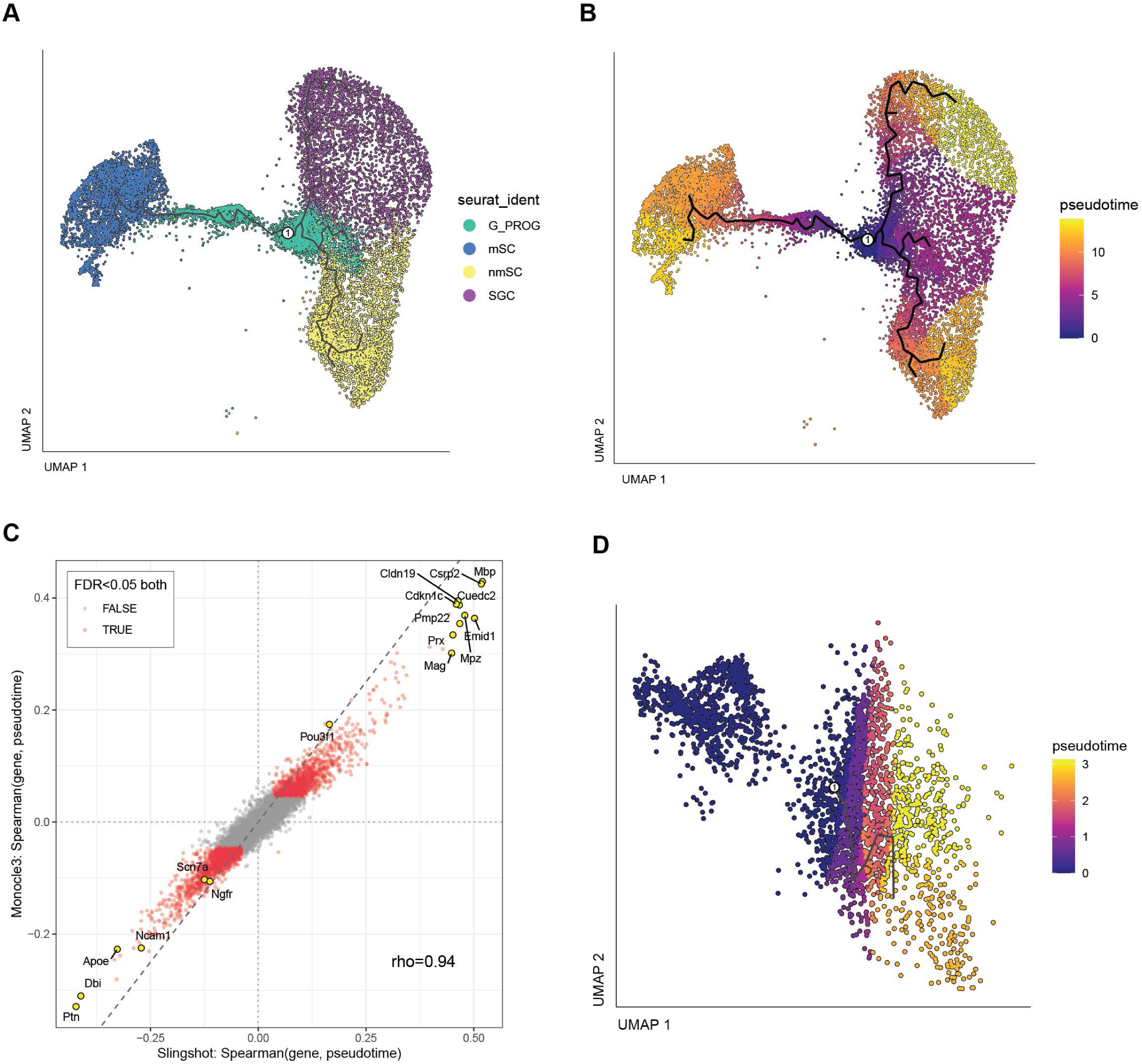
Robustness of the G_PROG trajectory inference confirmed by Monocle3 pseudotime, related to Figure 4. (**A**) Glia UMAP colored by major glial identity (G_PROG, nmSC, mSC, SGC). (**B**) Monocle3 pseudotime overlaid on the glia UMAP (principal graph in black), with the root selected as the cell with maximum Pou3f1 expression within the G_PROG cluster. (**C**) Per-gene Spearman correlation with pseudotime, comparing Slingshot and Monocle3 trajectory methods; overall correlation ρ=0.94 indicates robust agreement between the two methods. Genes significant (FDR<0.05) in both methods are highlighted; canonical lineage genes are labeled. (**D**) G_PROG-only Monocle3 pseudotime, restricted to G_PROG cells to highlight within-cluster ordering.

**Figure S14.**
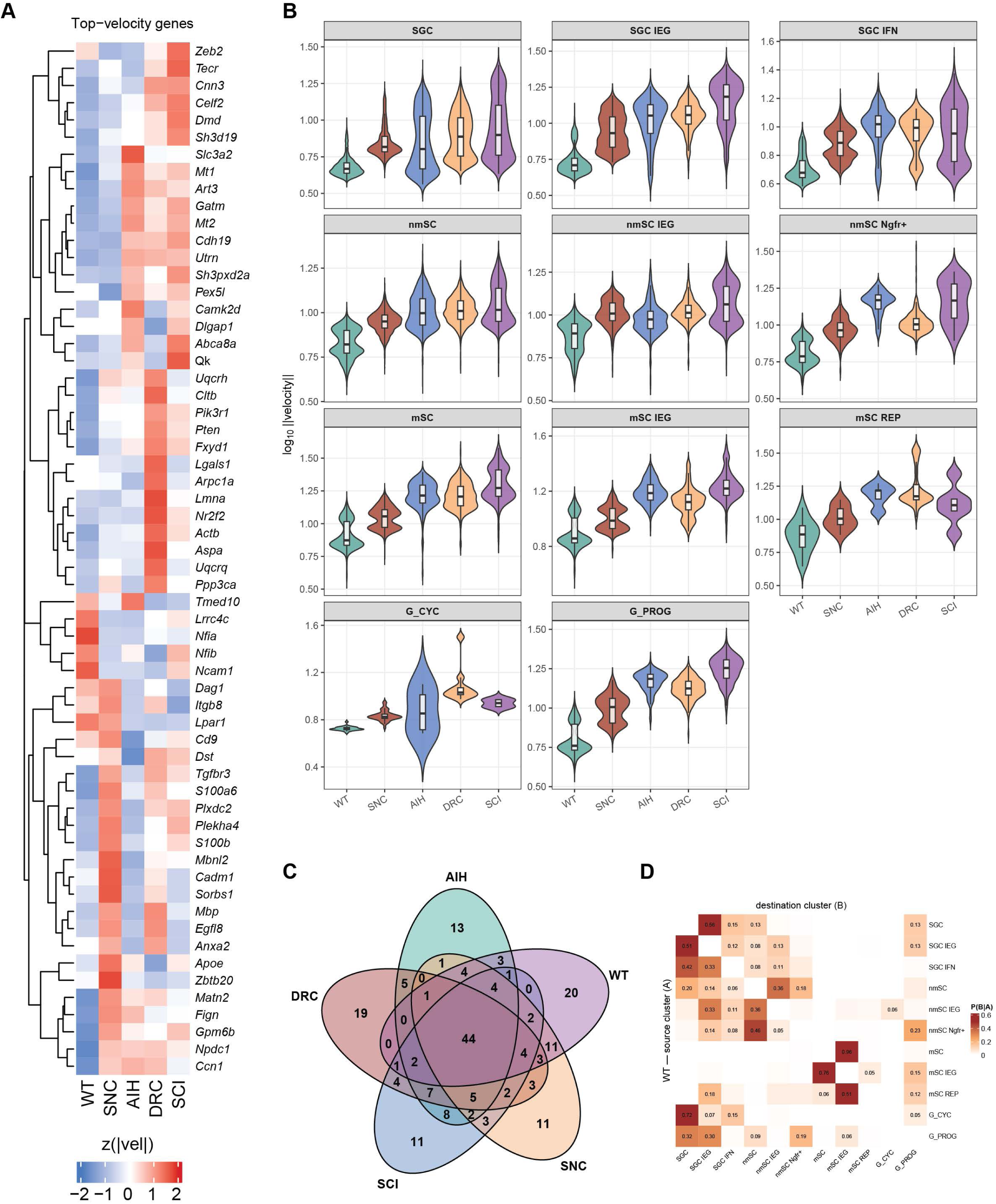
RNA velocity analysis confirms G_PROG trajectory directionality, related to Figure 4. (**A**) Heatmap of top genes driving spliced/unspliced velocity signal across glial Minor cell types. (**B**) Velocity magnitude violin plots split by condition (naïve, SNC, DRC, SCI, AIH) for each glial Minor cell type, demonstrating condition-specific induction of velocity-active gene programs. (**C**) Venn diagram showing the overlap of velocity-driving genes across the four injury conditions. (**D**) Per-condition heatmap of velocity transition probabilities between selected glial states, supporting directional transitions from G_PROG toward mature glial fates.

**Figure S15.**
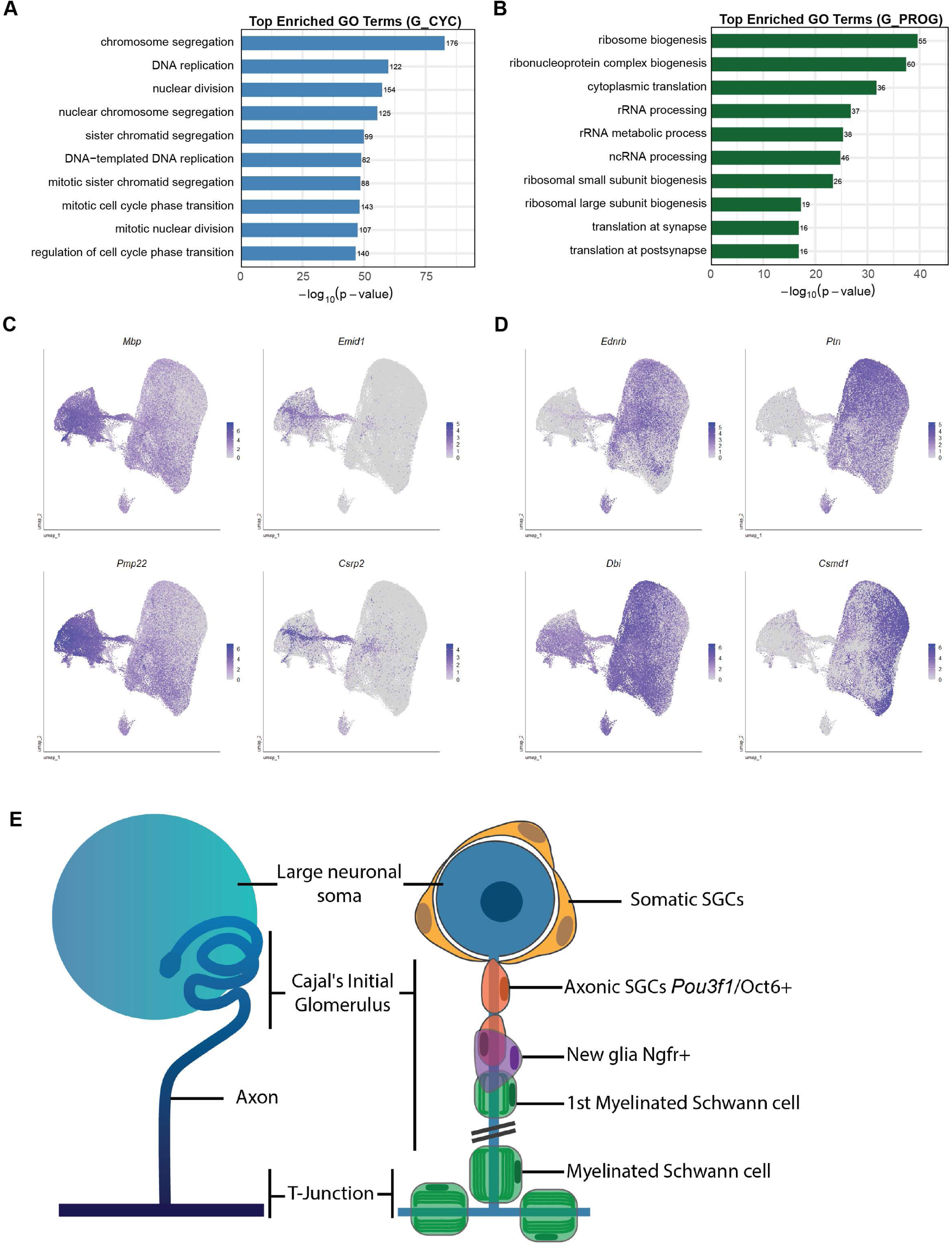
Characterization of progenitor-like glial cells in adult DRG, related to Figure 4. (**A-B**) Top 10 significant GO categories for significantly enriched in G_CYC population (**A**) and G_PROG population (**B**). Bar length indicates −log10(adjusted p-value). (**C-D**) UMAP feature plots showing expression of myelinating-associated genes; *Mbp, Emid1, Pmp22, Csrp2* (**C**) and non-myelinating-associated genes; *Ednrb, Ptn, Dbi, Csmd1* (**D**). (**E**) Schematic representation of a large neuron showing a 3D organization of the Cajal’s initial segment (left). On the right, a linear representation of the Cajal’s initial glomerulus and the organization of glial cells. The soma is covered with somatic SGCs, the proximal part of the Cajal’s initial segment is covered with axonal SGC before the first myelinating Schwann cell, which is covered by a new type of Ngfr+ glia.

**Figure S16.**
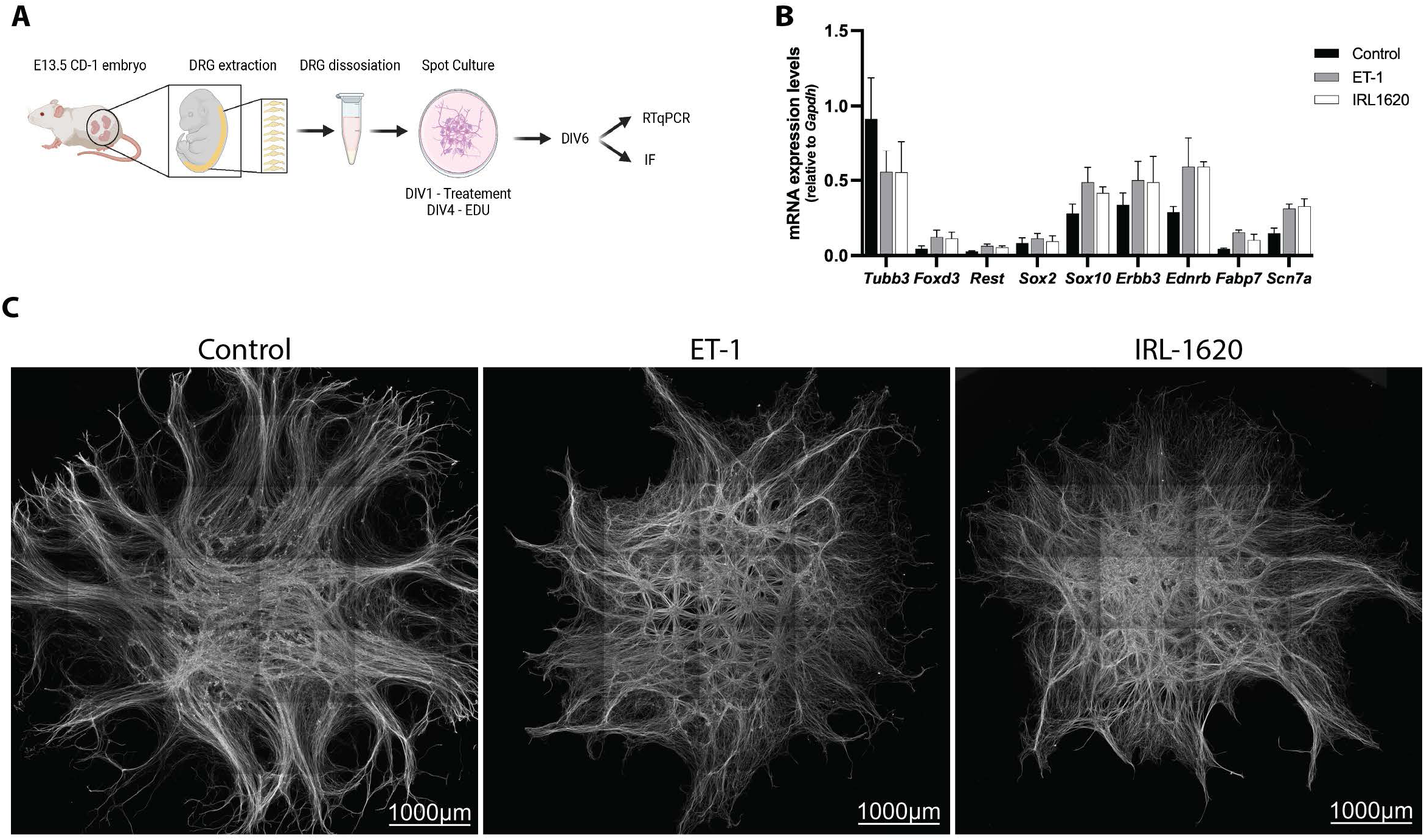
Endothelin signaling in embryonic DRG cultures, related to Figure 5. (**A**) Diagram of experimental design for eDRG culture studies. (**B**) Quantification of various cell markers using RTqPCR shows the expression of markers of neurons and glia and confirms the expression of *Ednrb* in eDRG spot culture treated or not. (**C**) Representative immunofluorescence images showing eDRG spot culture morphology using Tubb3 (gray) staining after 5 days of treatment with ET-1, IRL1620 compared to control. Scale bars: 1000 µm.

**Figure S17.**
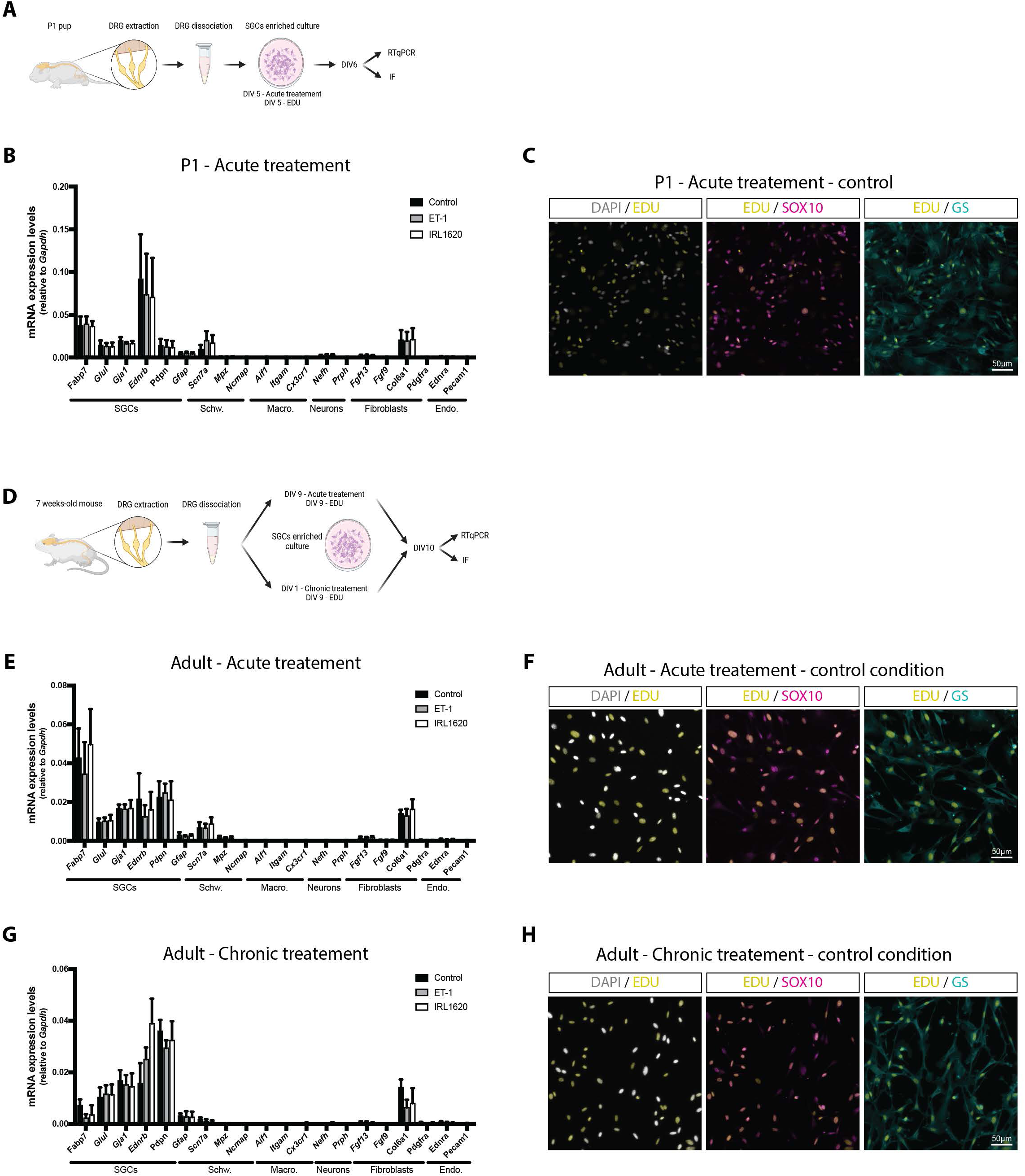
P1 and adult SGC culture characterization, related to Figure 6. **(A)** Diagram of experimental design for P1 SGCs culture studies. (**B**) Quantification of various cell markers using qPCR shows the higher expression of markers for SGCs over markers for Schwann cells (Schw.), macrophages (Macro.), neurons, fibroblasts (Fibro.) and endothelial cells (Endo.) treated or not. (**C**) Representative immunofluorescence images of control P1 SGCs culture showing EdU (yellow), Sox10 (magenta), GS (cyan), and Dapi (grey) staining. EdU+ nuclei co-localize with Sox10+ and GS+ cells. Scale bars: 50 µm. (**D**) Diagram of experimental design for adult SGCs culture studies. (**E, G**) Quantification of various cell markers using qPCR shows the higher expression of markers for SGCs over markers for Schwann cells (Schw.), macrophages (Macro.), neurons, fibroblasts (Fibro.) and endothelial cells (Endo.) under acute (**E**) or chronic (**G**) treatment and control conditions. (**F, H**) Representative immunofluorescence images of adult SGCs control culture showing EdU (yellow), Sox10 (magenta), GS (cyan), and Dapi (grey) staining. EdU^+^ nuclei co-localize with So×10^+^ and GS^+^ cells. Scale bars: 50 µm.

**Movie S1, related to Figure 1**.

3D UMAP visualizations of the integrated dataset (136,549 DRG cells; 90,972 SN cells) colored by hierarchical clustering level: class (left), major type (middle), and minor type (right). Example cluster annotations are shown below each UMAP to illustrate the hierarchical relationship between clustering levels.

**Movie S2, related to Figure 3**.

UMAP 3D visualization of 11 glial subpopulations identified by minor cell type clustering (SGC, satellite glial cell; SGC IFN, interferon-responsive SGC; SGC IEG, immediate early gene-expressing SGC; nmSC, non-myelinating Schwann cell; nmSC IEG, immediate early gene-expressing nmSC; nmSC Ngfr+, Ngfr-positive nmSC; mSC, myelinating Schwann cell; mSC IEG, immediate early gene-expressing mSC; mSC REP, repair mSC; G_CYC, mitotic glial cells; G_PROG, post-mitotic progenitors).

**Movie S3, related to Figure S15.**

Clarified mouse DRG stained with peripherin (red) enriched in neurons and Fabp7 (cyan) enriched in SGCs. The arrowheads point to the axons of neurons covered with SGCs.

**Table S1, related to Figure 1**

Summary of datasets integrated into the DRG/SN atlas. Each row represents an integrated batch containing one or more libraries from the indicated GEO accession(s). For accessions containing additional samples or conditions not listed here, only the indicated Treatment conditions were used. Study-level and cluster-level QC metrics are provided in subsequent sheets.

**Table S2, related to Figure 1**

Major cell type marker gene enrichment data. Differentially expressed marker genes for each of the 28 Major cell-type clusters (one sheet per cluster), computed via FindMarkers (Wilcoxon rank-sum test) against all other cells; columns as in Table S3. This table supersedes the major-marker spreadsheet from the original submission and was regenerated using the consolidated atlas in this revision.

**Table S3, related to Figure 3**

Glia Minor-level marker gene enrichment data. Differentially expressed marker genes for each of the 11 glia Minor cell-type clusters (one sheet per cluster), computed via FindMarkers (Wilcoxon rank-sum test) against all other glia cells; columns are gene, p_val, avg_log2FC, pct.1 (fraction of in-cluster cells expressing), pct.2 (fraction of all other cells expressing), and p_val_adj (Bonferroni-corrected).

## Key Resources Table

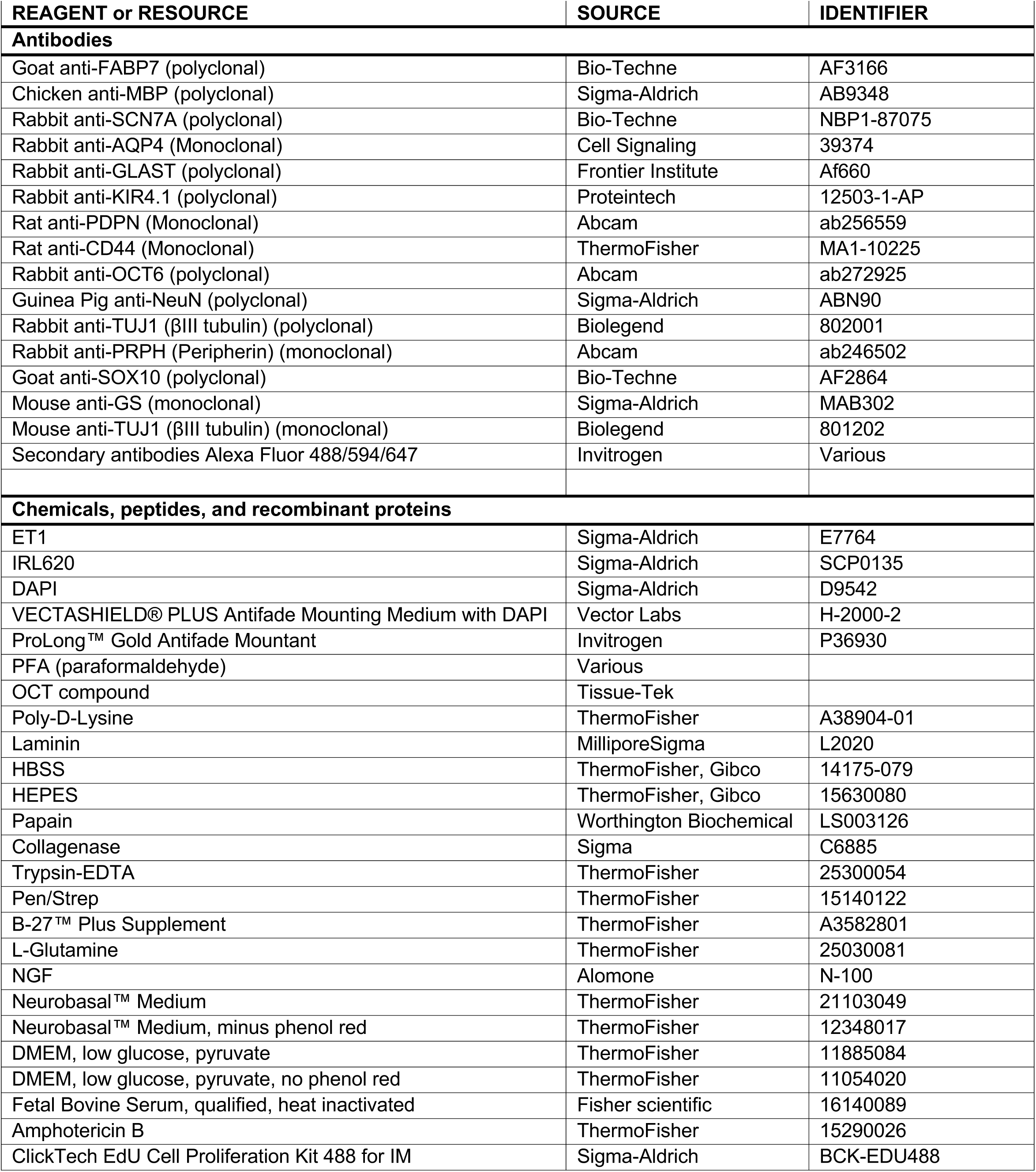
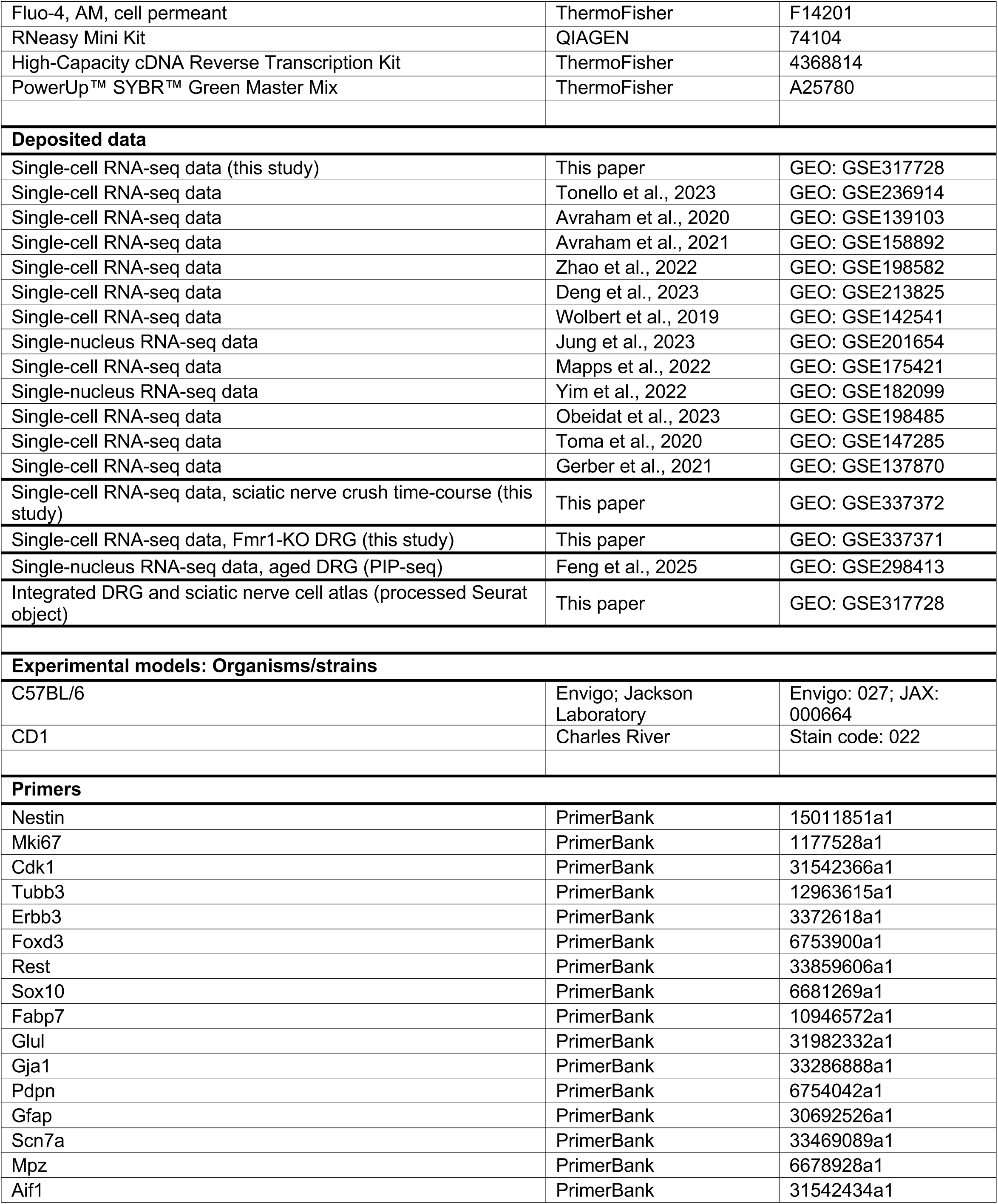
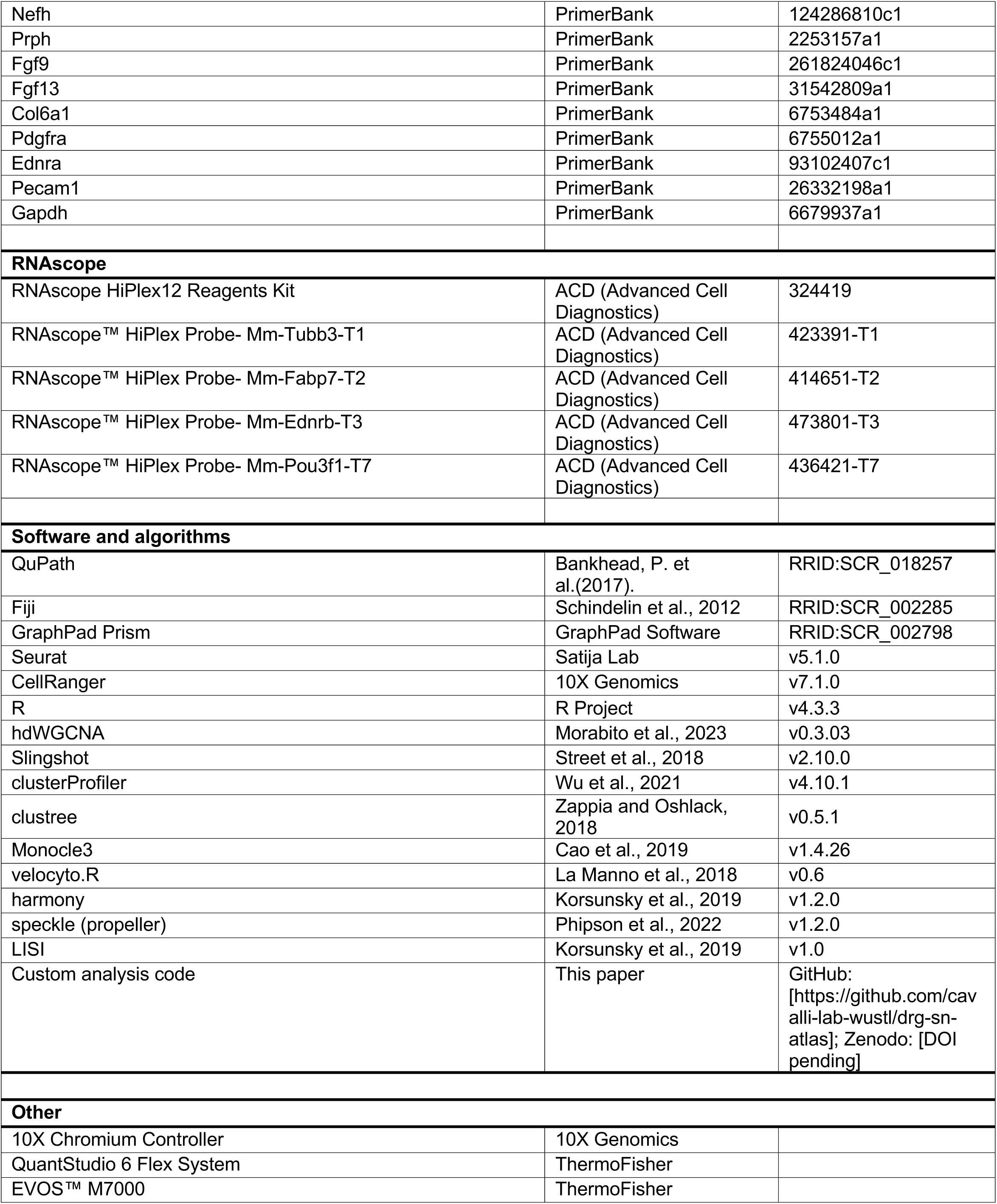
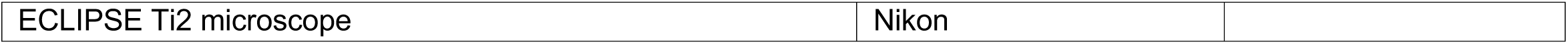

## Notes

### Competing Interest Statement

The authors have declared no competing interest.

### Summary of Updates

Results section updated to clarify; Figure 3 revised; Supplemental files updated.

